# A Chromosome-Scale Genome of *Nanomia septata* Reveals Extensive Rearrangement But No Clear Driver of the Unique Colony-Level Organization of Siphonophores

**DOI:** 10.1101/2025.05.01.651713

**Authors:** Namrata Ahuja, Darrin T. Schultz, Dalila Destanović, Samuel H. Church, Natasha Picciani, Catriona Munro, Koto Kon-Nanjo, Tetsuo Kon, Maciej K. Mańko, Wendy Shi, Steven H.D. Haddock, Oleg Simakov, Casey W Dunn

## Abstract

Siphonophores (Cnidaria:Hydrozoa) are pelagic colonial marine invertebrates with many highly specialized bodies (zooids) within a single colony. Their unique biology and ecological importance have made them of particular interest. Recent work revealed siphonophore genomes to be larger than in most other cnidarians. To investigate siphonophores’ genome biology and develop resources for future studies, we sequenced the genome of a single *Nanomia septata* to chromosome scale. The haploid genome is 1.5GB across 8 chromosomes, a reduction relative to the 15 chromosomes seen in closely related hydrozoan genomes, and is highly rearranged, consistent with multiple mixing events. Genome expansion occurred through intergenic repeat expansion, with protein-coding genes shorter than in most cnidarians. We found no genomic features clearly associated with siphonophores’ exceptional colony-level complexity. Gene families that play critical roles in cnidarian development have not expanded, and gene proximity was not generally correlated to their expression across zooids, except in male gonophores. To contextualize these observations, we genome sequenced 20 additional *Nanomia* specimens across the globe and mapped them to our chromosome-scale reference. Population genomic analyses support three previously recognized species of *Nanomia*, and at least one additional undescribed species. Overlapping geographic distribution of some *Nanomia* species suggest reproductive isolation in sympatry. Phylogenetic analyses of genome size indicate *Nanomia septata* and *Nanomia cara* have similarly large genomes between 1.5-1.7GB, while *Nanomia bijuga* and an undescribed species show a secondary reduction to 0.7GB. These results highlight how genomic factors have shaped colony organization and genome diversity within *Nanomia*.

## Introduction

Siphonophores (Cnidaria:Hydrozoa) are animals with complex colony-level body plans (Dunn and Wagner 2006; Mapstone 2014). Though each siphonophore colony grows from a single embryo, they asexually produce many specialized bodies, or zooids, each with distinct functions and morphologies (Mackie 1964; Mackie et al. 1988; Dunn and Wagner 2006). These bodies share resources through a common gastrovascular cavity and are highly physiologically integrated (Mackie et al. 1988; Dunn and Wagner 2006). This functional specialization of bodies within a colony has long made siphonophores of central interest to questions about the evolution of new levels of biological organization, analogous to the transition from unicellularity to multicellularity or to the evolution of functional specialization of cells within a body (Haeckel 1869; Mackie 1963; Beklemishev 1969).

*Nanomia septata* (Figure 1A), has emerged as a particularly useful model species for the study of these unique features of siphonophore biology. This species is abundant in coastal waters of the northeast Pacific, where it can be reliably collected (Hosia et al. 2024). Studies have described the budding process by which the colony grows (Dunn and Wagner 2006), assessed differential gene expression across their zooids (Munro et al. 2022), examined the spatial patterning of i-cell genes within a colony (Siebert et al. 2015), and characterized some of the cell types present in the colony (Church et al. 2015). The Monterey Bay Aquarium has kept *Nanomia septata* alive for long periods of time in culture (Patry et al. 2020). Further studies using *Nanomia septata* to advance general questions about siphonophore biology, including their unique colony-level growth and organization, would benefit greatly from a chromosome scale genome.

**Figure 1.**
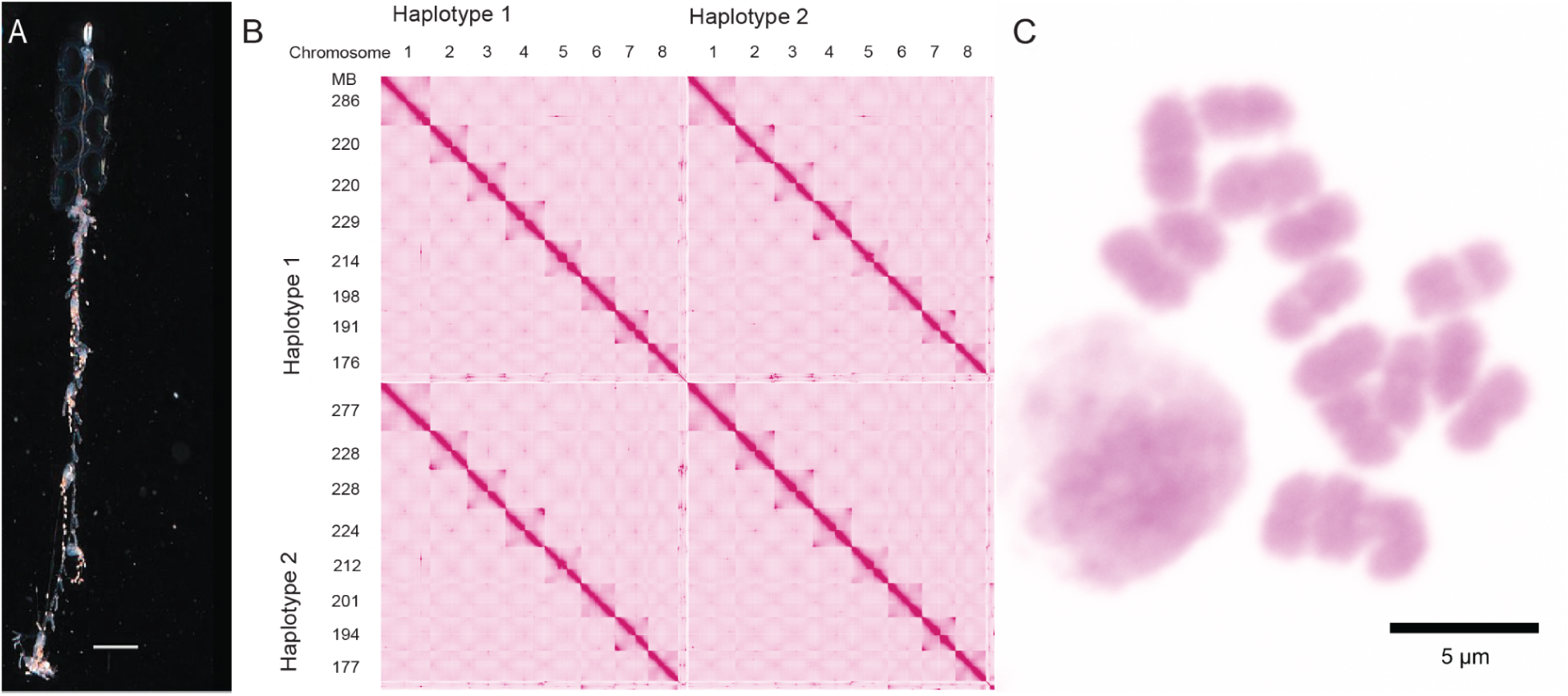
Genome of *Nanomia septata*. A. *Nanomia septata*. Scale bar is 1 cm. Photo credit to C. Munro. B. Omni-C Heatmap of genome assembly. Haplotype 1 is on the upper left, and haplotype 2 is on the lower right with the upper right and lower left squares representing the overlap between both haplotypes, Each chromosome is numbered 1-8 for each haplotype along the top row. Numbers on the left side give the approximate length of each chromosome in MB. C. *Nanomia septata* karyotype showing 2n=16. A stained nucleus is present in the bottom left.

Multiple cnidarian genomes have been sequenced to chromosome-scale in recent years. They tend to be relatively small for animal genomes, at 300-700MB (Putnam et al. 2007; Chapman et al. 2010; Leclère et al. 2019; Kon-Nanjo et al. 2023; Kon-Nanjo et al. 2024), though larger cnidarian genomes have been found (Law et al. 2023; Church et al. 2024). Within Hydrozoa, siphonophores are most closely related to Filifera III + Filifera IV (Bentlage and Collins 2021), a group that includes several well studied species including *Hydractinia symbiolongicarpus*, *Hydractinia echinata*, and *Podocoryna carnea*. Macrosynteny is highly conserved across many non-siphonophore cnidarians (Simakov et al. 2022)*. Hydra vulgaris* and *Hydractinia symbiolongicarpus* form a paraphyletic grade with respect to siphonophores, with *Hydractinia symbiolongicarpus* being the closer relative to siphonophores (Bentlage and Collins 2021). High quality chromosome scale genome assemblies reveal that *Hydra vulgaris* and *Hydractinia symbiolongicarpus* have strikingly similar genome organizations, each with 15 chromosomes, similar genome content, and sizes of 912 MB and 483 MB, respectively (Chapman et al. 2010; Simakov et al. 2022; Kon-Nanjo et al. 2023). The conserved features of these two species, therefore, provide a clear picture of the ancestral genome structure, likely with 15 haploid chromosomes and a small genome, from which the genomes of siphonophores evolved.

It was surprising, then, when a new study on genome sizes of 35 siphonophores showed that many siphonophore genomes are larger than most other cnidarians (Ahuja et al. 2024)*. Nanomia* was particularly interesting since *Nanomia septata* has a genome size around 1.4GB, larger than typical of other cnidarians but within the range of siphonophore genome sizes, while *Nanomia bijuga* has a much smaller genome of only about 0.7GB (Ahuja et al. 2024). This previous work found that the *Nanomia septata* genome is about 63% repetitive (Ahuja et al. 2024), partially explaining the large size, though the types of repeats present were unknown. The genome differences between these two *Nanomia* species offers an opportunity to expand our understanding of population differences and diversity at the genomic level.

Only two species of *Nanomia*, *Nanomia bijuga* globally and *Nanomia cara* in the North Atlantic, were recognized as valid for most of the past century (Bargmann and Totton 1965). A new study based on morphology and sequences of two mitochondrial genes made critical progress toward understanding global *Nanomia* diversity (Hosia et al. 2024). It presented strong support for a distinct species in the North Pacific, which the authors describe as *Nanomia septata* (Mapstone 2014; Hosia et al. 2024). Before 2024, all specimens of *Nanomia septata* had been referred to as *Nanomia bijuga* in the literature (though not all specimens referred to as *Nanomia bijuga* are *Nanomia septata*). This previous study also found several *Nanomia* specimens that are likely from additional undescribed species (Hosia et al. 2024). Newer and improved methods and tools are well suited to advancing our rapidly changing picture of diversity of open ocean animal systems, including on *Nanomia*. Questions on genome biology and diversity are critical to using *Nanomia* as a model organism for the study of siphonophore biology, as well as addressing general questions of broad interest about the structure of animals in the open ocean.

## Results

### Genome Description

#### Genome structure and annotation

We present a highly contiguous, complete chromosome-scale diploid assembly of a single *Nanomia septata* colony collected near Monterey, California (Figure 1B). Each of the two haplotypes have 8 chromosomes and a BUSCO score of ∼92% (Table 1 and Figure 1). A karyotype confirmed there are 2n=16 chromosomes (Figure 1C). We designate each chromosome with a number 1-8. Haplotype 1 has 96.7% of the bases in 8 chromosomes, and haplotype 2 has 96.9% of bases in 8 chromosomes. The haplotypes are very similar to each other, with haplotype 2 having a slightly higher completeness and assembly quality (Table 1).

**Table 1.** *Nanomia septata* genome assembly statistics.

| Description | Haplotype 1 (alternate) | Haplotype 2 (primary) |
| --- | --- | --- |
| Length | 1794626896 | 1796533731 |
| Contigs | 5979.0 | 4964 |
| Contig N50 | 888919 | 918025 |
| Scaffolds | 607 | 468 |
| Scaffold N50 | 220372998.0 | 223932061.0 |
| BUSCO complete | 91.8 | 92.2 |
| BUSCO single copy | 85.4 | 85.4 |
| BUSCO duplicated | 6.4 | 6.8 |
| BUSCO fragmented | 2.8 | 2.8 |

Haplotype 2 also had fewer gaps than the alternate, and contained more contigs within the chromosome-scale scaffolds, resulting in fewer orphan scaffolds. Thus, we chose to use haplotype 2 for subsequent analyses, including annotation (Table 2).

**Table 2.** *Nanomia septata* annotation of haplotype 2 (primary).

|  |  |
| --- | --- |
| Number of genes | 26,323 |
| Total gene length | 110.4 MB |
| Total length of introns | 71.3 MB |
| Total length of exons | 37.4 MB |
| Intron ratio | 2.3027 |

We manually curated the *Homeobox* and *Wnt* gene annotation, given their importance in cnidarian development (body and axis patterning) (Sinigaglia et al. 2013; He et al. 2018; Technau and Genikhovich 2018; Holstein 2022) and potential relevance to the unique colony-level growth of siphonophores . The original annotation had 38 of these genes and we found an additional 22 more using the iso-sequencing of *Nanomia septata.* In total, *Nanomia septata* has 60 homeobox genes, split across 8 classes of genes (Supplementary Table 1). In comparison, other hydrozoans have on the order of 60-80 homeobox genes (Supplementary Table 1). More specifically, *Nanomia septata* has 17 Antp genes, including both Hox-like (HOXL) and NK-like (NKL) members. Among HoxL genes, we identified one Hox gene (Cnox-1, located on chromosome 4), an unclassified Hox-like gene, and the two paraHox genes Gsx (Cnox-2) and Cdx (Cnox-4). We found 13 Nk-like genes, including Nk-1, Nk-2 (three copies), and Nk-6 genes (Supplementary Figure 1). In addition, it has eight Wnt genes identified as *Wnt* 1, 2, 3, 5 (two copies), 6, 7 and A (Supplementary Figure 2). These have all been added so that in total, the annotation of haplotype 2 has 26,323 protein-coding genes (the first version had 26,301 genes).

The total length of introns in the final assembly is 71.3Mb and total length of exons is 37.4Mb, with a mean intron ratio (Glick et al. 2024) of 2.3027 (Table 2).

#### Differential gene expression across zooids

A previous bulk RNA-seq study assessed the differential expression of genes across different types of developing and mature zooids in 7 species using short read transcriptomes assembled with TRINITY as references (Munro et al. 2022). We re-mapped the *Nanomia* RNA-seq reads used in that study to our new genome and annotation. The results from the original paper (Munro et al. 2022) are robust to the improved gene references. For example, female gonodendron and nectophores group together (Supplementary Figure 3). We included the male gonodendron dataset that was excluded in the original paper and found that it also closely groups with the female gonodendron and nectophores (Supplementary Figure 3), consistent with all these zooids being highly modified medusae that are retained within the colony. The pneumatophore, a specialized structure for buoyancy that is not a zooid, clusters closely with both gonodendra and nectophores (Supplementary Figure 3). Lastly, gastrozooids and palpons, which are both types of specialized polyps, have more similar expression to each other than to other zooids.

These re-mapped data also provide an updated list of genes with zooid-specific expression. We find that there are a total of 3,816 genes that have significant differential expression among zooid tissues and stages (Supplementary Table 2). Of these, 3,782 genes have expression specific to a single zooid type, mostly with increased expression in either mature female gonodendra or developing nectophores (Supplementary Table 2). Similar to the original study (Munro et al. 2022), genes with elevated expression in gastrozooids include many with functions related to digestion. With our new results, we also identify GOs (gene ontologies) for toxins that are abundant in female gonodendra (GO:0019835, GO:0030286, GO:0032991), nectophores (GO:0030286), palpons (GO:0004866) and the pneumatophore (GO:0004866) (Supplementary Table 2).

The remaining 34 out of 3,816 total genes have differential expression that is shared across multiple zooid types (Supplementary Table 3). Unsurprisingly, developing nectophores have shared differential expression with the mature pneumatophore and the mature female gonodendra, all of which cluster together in the PCA (Supplementary Table 3 and Supplementary Figure 3). These shared differentially expressed genes have GOs for developmental processes such as morphogenesis of a branching structure and morphogenesis of epithelium. Specifically, TAGLN3, an actin gene, is shared and upregulated between the female gonodendra and pneumatophore, and UNC45A, a muscle development gene, is shared and upregulated between the female gonodendra and nectophore (Supplementary Table 3).

Fox/forkhead genes have elevated expression in female gonodendra and developing nectophores, consistent with their role in medusa development observed in other cnidarians (Chevalier et al. 2006). This updated list of differentially expressed genes in zooids will be useful for pursuing future in-depth zooid studies, including spatial expression experiments and single cell sequencing (Supplementary Table 2).

Additionally, we assessed the relationship between physical proximity of genes and the covariance of their expression across zooid types. We found multiple regions in the genome enriched for genes with correlated expression across zooid types (Supplementary Figure 4). In all cases, these genes had elevated expression specific to male gonophores. These genes included spermatogenic genes (Supplementary Figure 4).

#### Repeat Analysis

63.7% of the genome is composed of repetitive elements (Figure 2; Supplementary Table 4,5). This is consistent with previous estimates based on k-mers (Ahuja et al. 2024). Notably, the proportion of DNA elements was high, with *TcMar-Tc1*, *Kolobok-T2*, *Maverick*, *PiggyBac*, and *hAT-Charlie* being abundant (Figure 2B). In addition, many other TEs are also abundant in the genome (Figure 2B,C; Supplementary Figure 5C, 6; Supplementary Table S4). Furthermore, when examining the substitution levels of subfamilies of each transposon, we found a large number of subfamilies with low divergences (Figure 2A). This suggests that the *Nanomia septata* genome contains many young/active transposons. Additionally, other species belonging to *Nanomia* have been reported to exhibit a wide range of genome sizes (Ahuja et al. 2024).

**Figure 2.**
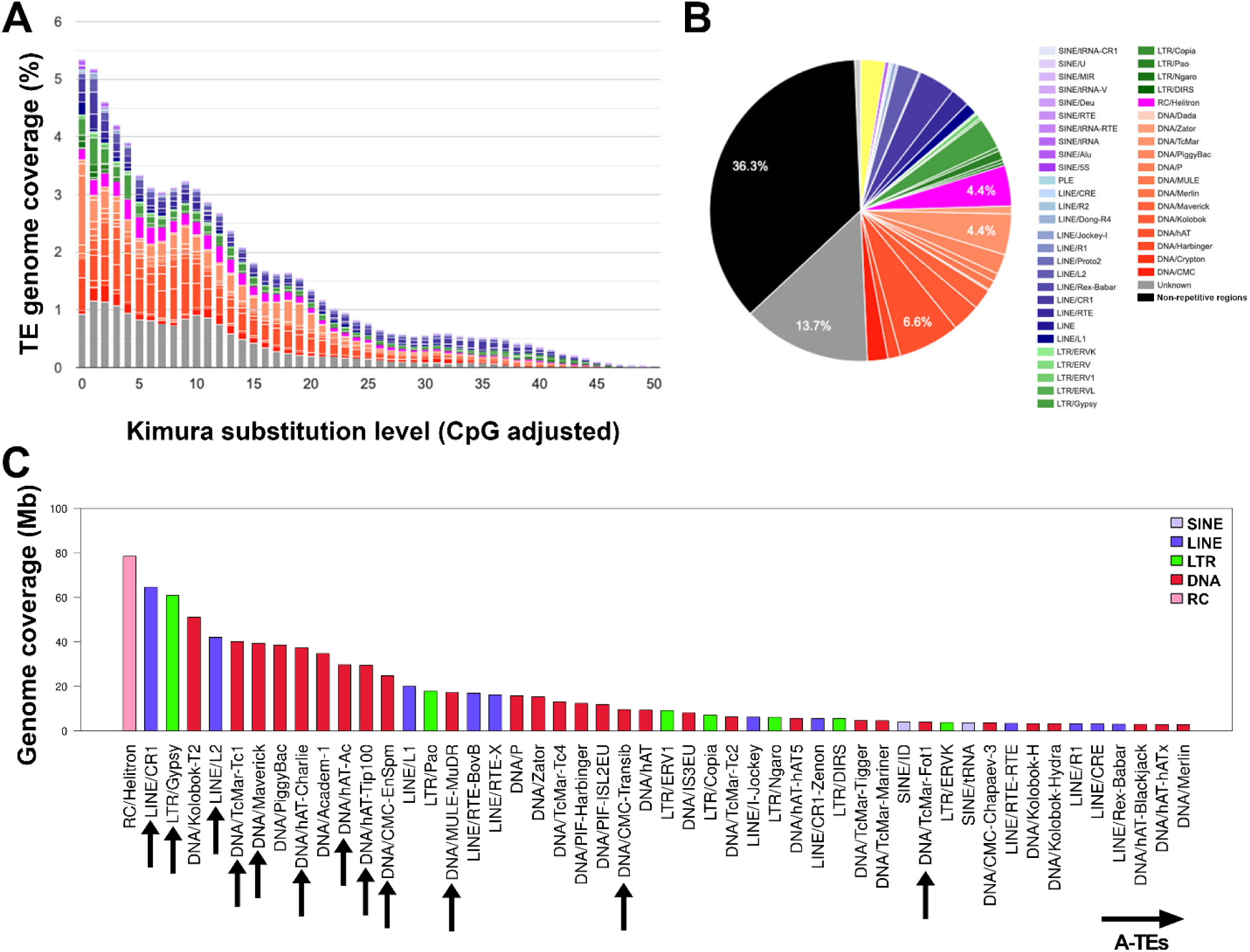
Repeat landscape of the *Nanomia septata* genome. A. Genome coverages and sequence divergences of TE families. The x-axis depicts the degree of Kimura substitution within repeat elements compared to consensus sequences, serving as an indicator of their relative age. The y-axis shows the genome coverage of each repeat family throughout the genome. Repeats with ancient activity are located towards the graph’s right, while those with more recent activity are found on the left side. B. Pie chart representing proportional representation of each TE family. Note that 36.3% of the genome consists of non-repetitive regions, while the remaining 63.7% are repetitive elements. C. Absolute genome coverage of top 50 TEs. Arrows indicate the eukaryotic Core-TEs (Kon-Nanjo et al. 2024). The eukaryotic Core-TEs collectively occupy 28.1% (505 Mb) of the nanomia genome.

Previously, Kon-Nanjo et al. (2024) identified 14 highly conserved active TE families (A-TEs), mainly composed of DNA elements (hAT-Ac, hAT-hATm, hAT-Tip100, hAT-Charlie, TcMar-Tc1, TcMar-Fot1, CMC-Transib, CMC-EnSpm, Sola-2, Maverick, and MULE-MuDR), that are found across a wide range of eukaryotic genomes. In the *Nanomia septata* genome, these A-TEs collectively occupy 22.3% (401 Mb) of the genome (Figure 2C; Supplementary Figure 5). We also compared nine species of metazoans including two bilaterians (*Pecten maximus* and *Branchiostoma floridae*), seven cnidarians (*Nematostella vectensis*, *Acropora millepora*, *Rhopilema esculentum*, *N. septata*, *Hydractinia symbiolongicarpus*, *Hydra viridissima*, and *Hydra vulgaris*). *Nanomia septata* exhibited greater genome coverage by a wider variety of TE families than any other analyzed animal, indicating a multi-TE family expansion (Cluster #1 in Supplementary Figure 5C, 6). Additionally, 13.7% of repeats across the entire genome could not be classified, suggesting that they may be specific to *Nanomia septata* or siphonophores more broadly.

### Comparative cnidarian genomics

Many features of synteny are conserved across more than 500M years of cnidarian evolution, but are then lost in siphonophora (Figure 3A). This indicates that siphonophore genomes have undergone rapid change relative to other cnidarian genomes, and are quite distinct in many features. These changes can be partitioned between evolution along the siphonophore stem branch (i.e, after siphonophores diverged from other cnidarians but before the most recent common siphonophore ancestor), and later genome evolution within siphonophores. Because *Hydra vulgaris* and *Hydractinia symbiolongicarpus* form a paraphyletic group with respect to siphonophores (Bentlage and Collins 2021), their many shared features provide extensive information about the state at the start of the siphonophore stem branch (Figure 3A). These shared ancestral features include 15 chromosomes with highly conserved synteny (Figure 3A), and a typically small cnidarian genome size. Given the similarities between these two species, we make comparisons to *Hydractinia symbiolongicarpus* as a proxy for the ancestral state.

**Figure 3.**
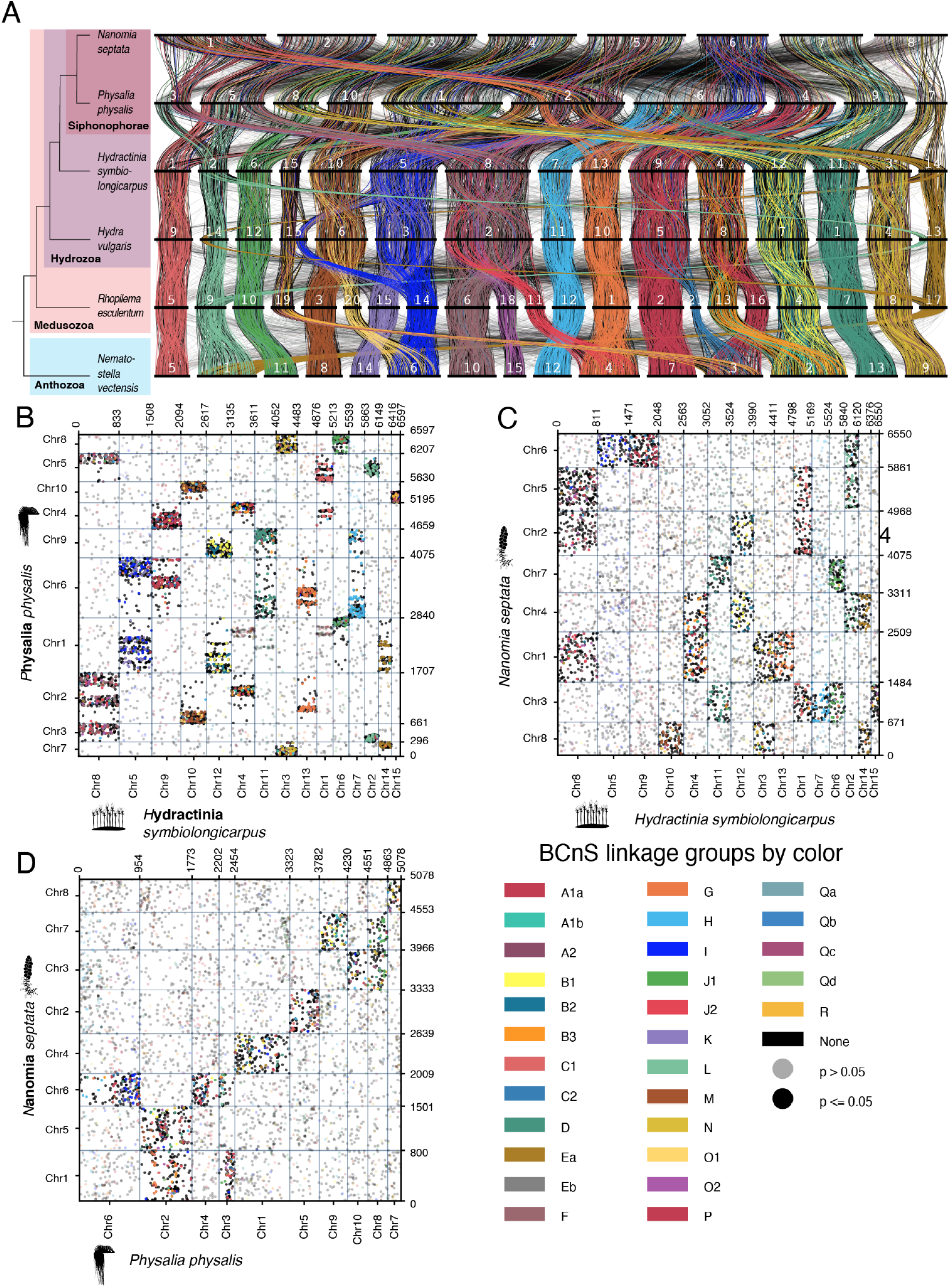
*Nanomia septata* chromosomes are highly rearranged. A. Ribbon plot of *Nanomia septata* vs other cnidarians. B-D. Oxford dot plots (ODP) of pairwise hydrozoan comparisons. B. ODP of *Hydractinia symbiolongicarpus* chromosomes vs *Nanomia septata* chromosomes. C. ODP of *Hydractinia symbiolongicarpus* chromosomes vs. *Physalia physalis* chromosomes. D. ODP of *Physalia physalis* chromosomes vs *Nanomia septata* chromosomes. Chromosomes are below the dot plots and are colored along with gray and black representing p values.

We compared the *Hydractinia symbiolongicarpus* and two siphonophore genomes, *Physalia physalis* and *Nanomia septata* using oxford dot plots (Figure 3B-D). A chromosome scale assembly of the siphonophore *Physalia physalis* was recently produced for a population analysis of *Physalia* (Church et al. 2024) and is used here. Comparison among these three species indicate that there is not a 1:1 correspondence between *Hydractina symbiolongicarpus* chromosomes (and other cnidarians) and any chromosomes in either siphonophore species.

The *Physalia physalis* genome is more similar than *Nanomia septata* to *Hydractinia symbiolongicarpus*, indicating that the genome of *Nanomia septata* has undergone more extensive reorganization (Figure 3). There is stronger statistical support for chromosomes between *Hydractina symbiolongicarpus* and *Physalia physalis* (Supplementary Table 6: Fig3_Chr_Values_PanelB), while support values for chromosomes shared between *Hydractina symbiolongicarpus* and *Nanomia septata* are marginal (Supplementary Table 6: Fig3_Chr_Values_PanelC). There are discrete areas of homology between pieces of *Hydractinia symbiolongicarpus* chromosomes and pieces of *Physalia physalis* chromosomes, such as chromosome 15 in *Hydractinia symbiolongicarpus* which corresponds to chromosome 10 in *Physalia physalis* (Figure 3B). In contrast, we observe scrambling of orthologs across the entire *Nanomia septata* genome, suggesting multiple translocations (Figure 3C). To understand the degree of scrambling of the previously described ancestral linkage groups (ALGs; bilaterian-cnidarian-sponge and unicellular-metazoan linkage groups), we visualize these ALGs in *Hydractinia symbiolongicarpus*, *Physalia physalis*, *Nanomia septata* (Supplementary Figure 7) which provides additional support for more extensive genome reorganization in *Nanomia septata*. All the supporting values for the pairwise analyses above are contained in Supplemental Table 4.

Next, we looked at the comparison between the *Nanomia septata* and *Physalia physalis* genomes (Figure 3). Features shared uniquely between *Physalia physalis* and *Nanomia septata* reveal information about the most recent common ancestor of siphonophores (ie, at the end of the siphonophore stem branch). Differences between *Physalia physalis and Nanomia septata* reveal evolutionary changes within *Siphonophora.* There have been many fusion events but there are still a few detectable ones. These suggest that the reduced karyotype in *Nanomia septata* (n=8) and *Physalia physalia* (n=10) could have arisen by some fusions of the ancestral homologs of *Physalia physalis* chromosomes 2 and 3 to form LG 1, chromosomes 8 and 10 to form LG 3, and fusions of chromosome 6 and 4 to form LG 6 in *Nanomia septata* (Figure 3D). However, these are what are detectable, meaning more translocations and mixing events are present that are hard to pin down with only 2 siphonophore genomes.

### Genomic diversity in *Nanomia*

To assess the broader context of diversity and population structure in *Nanomia*, we performed whole genome short read sequencing on 24 specimens collected over 19 years from the Atlantic Ocean (Virginia and Rhode Island), Pacific Ocean (Washington, California, Gulf of California, and Hawai’i), and Mediterranean Sea (Villefranche-sur-mer). Phylogenetic analyses of CO1 (Supplementary Figure 8) indicate that these specimens belong to *Nanomia septata* (7 samples from California, 3 from Washington), *Nanomia cara* (1 sample from Rhode Island), *Nanomia bijuga* (3 from Rhode Island, 2 from Hawai’i, 1 from Villefranche-sur-mer, 3 from Gulf of California), and one undescribed species from Hawai’i consistent with *Nanomia* sp. 1 from Hosia et al (2024).

We assembled mitochondrial genomes from all sampled *Nanomia* and found them all to have the same conserved gene order recently reported for *Nanomia septata* and *Nanomia bijuga* (Ahuja et al. 2024). The phylogenetic relationships of the full-length mitochondrial genomes of *Nanomia* (Figure 4A) are consistent with the relationships based on 16S and CO1 reported by Hosia et al (2024), with improved support (bootstrap values at all internal nodes 100), and with our CO1 analyses (Supplementary Figure 8). *Nanomia cara* is sister to all other *Nanomia,* and *Nanomia septata* is sister to a clade of *Nanomia bijuga* and *Nanomia* sp. 1 (Figure 4).

**Figure 4.**
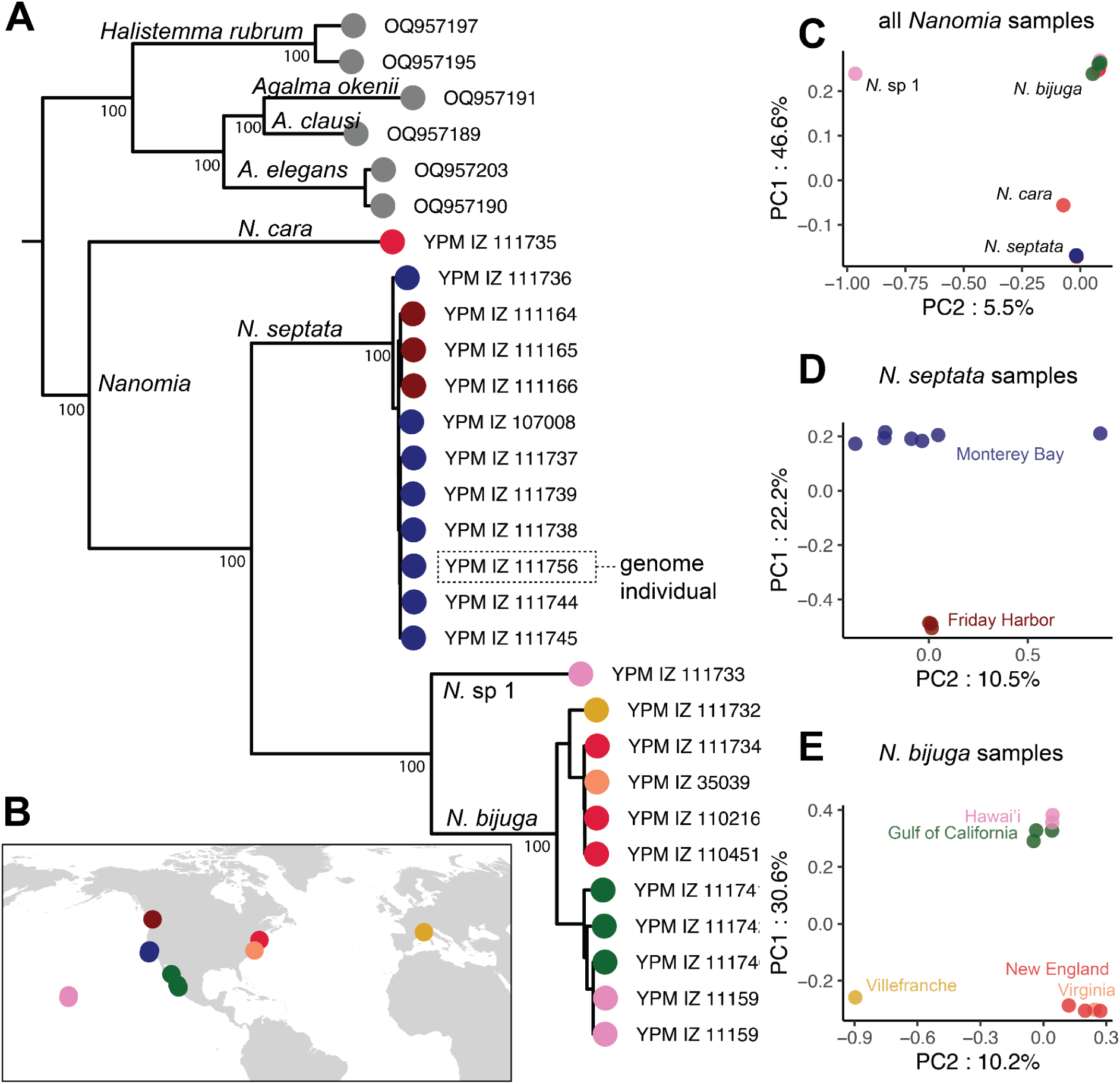
Population genomics of *Nanomia*. A. Phylogeny based on complete mitochondrial genome. B. Map of collecting locations, which are coded by color. Color in other panels correspond to locations here. C. PCA of all *Nanomia* specimens newly sequenced in this study. D. PCA of *Nanomia septata* specimens only. E. PCA of *Nanomia bijuga* samples only. All PCAs use the Nanomia *septata* reference genome. Analyses with species specific transcriptome references are shown in Supplemental Figure 8.

We also used k-mer distributions (Supplementary Figure 9) to confirm and extend recent observations of genome sizes of *Nanomia septata* and *Nanomia bijuga* (Ahuja et al. 2024). All samples of *Nanomia septata* have genome sizes of 1.5GB and have similar repeat and heterozygosity statistics consistent with the single individual analyzed previously (Ahuja et al. 2024). The single sample of *N. cara* had a 1.7GB genome, providing critical new information on a species for which no previous genome statistics were available. K-mer analyses of all *N. bijuga* samples, as well as the specimen of *Nanomia* sp 1 show that these species have much smaller genomes of about 0.7GB and a lower fraction of repetitive elements. Taken together, these results indicate that there are two *Nanomia* species, *Nanomia septata* and *Nanomia cara*, with large genomes of 1.5-1.7GB, and two, *Nanomia bijuga* and *Nanomia* sp. 1 with much smaller genomes that are about half the size at 0.7GB. In combination with the phylogenetic results that show *N. septata* sister to *N. bijuga* and *N.* sp. 1 and *N. cara* sister to all of these species (Figure 4), these analyses suggest that the most recent common ancestor of *Nanomia* had a genome size of about 1.7GB, and that there was then a large reduction in genome size along the branch that gave rise to *Nanomia bijuga* + *Nanomia* sp. 1 (Figure 4).

Combined population genomic analyses (see Methods), based on mapping each genome to the chromosome scale reference genome for *Nanomia septata*, clearly differentiate the species identified above (Figure 4C). There is also strong geographic structure to population diversity within species. Within *Nanomia septata* (Figure 4D), samples collected near Friday Harbor, Washington are distinct from those collected near Monterey, California. Within *Nanomia bijuga* (Figure 4E), we observe a separation between Pacific specimens and other specimens on PC1, and between the Mediterranean specimen and other specimens on PC2 in the graph (Figure 4E).

Reads from *Nanomia septata* map well to the *Nanomia septata* reference genome (85%–93% properly paired), but reads from *Nanomia cara* (44%), *Nanomia bijuga* (22–67%), and *Nanomia* sp. 1 (44%) map poorly (Figure 4). Previous work in *Physalia* found that results using transcriptome and genome references are highly similar (Church et al. 2024). To test whether results were robust to choice of reference sequence in *Nanomia*, we generated transcriptomes (iso-seq) for *N. bijuga* and *N. septata* and used these as a reference for mapping reads from those species (Supplementary Figure 10). The PCAs of variation using species-specific transcriptome references are largely consistent with those derived when mapping to the reference genome (Supplementary Figure 10).

*Nanomia* species have distinct but overlapping ranges (Figure 4 and Supplementary Figure 10C). For example, *Nanomia septata* is present across the North Pacific, from Japan, Alaska, and California. Likewise, *Nanomia bijuga* is found in parts of the Pacific, including Hawai’i, Japan, and Mexico, but also in the Atlantic and Mediterranean. Japan is a spot of overlap from *Nanomia septata* and *Nanomia bijuga* and *N*anomia sp. 1 (Hosia et al 2024).

Additionally, *Nanomia* sp. 1 and *Nanomia bijuga* were both collected in Hawai’i, and *Nanomia cara* and *Nanomia bijuga* were both collected in Rhode Island.

## Discussion

A recent study found that most siphonophores, including *Nanomia septata*, have very large genomes relative to other cnidarians (Ahuja et al. 2024). The chromosome-scale assembly of *Nanomia septata* presented here (Figure 1B) extends these findings to reveal that the genome of this siphonophore is highly derived in multiple respects, not just size. It has expanded repetitive elements in intergenic regions (Figure 2), as expected of a larger genome, but intragenic elements have contracted. There has also been a reduction from the 15 chromosomes seen in non-siphonophore hydrozoans to 8 chromosomes in *Nanomia septata*, a process that included extensive translocations and fusion-and-mixing, which has led to a highly reorganized genome in this species. Synteny analyses (Figure 3) with multiple cnidarian genomes, including an unannotated chromosome-scale reference genome from a population genomic analysis of the siphonophore *Physalia physalis* (Church et al. 2024), reveal that there has been extensive genome reorganization along the stem branch that gave rise to the most recent common ancestor of siphonophores and then within siphonophores. Extensive genomic rearrangements could be a result of rapid chromosome evolution, as seen in other metazoans including clitellates, tunicates and now siphonophores (Lewin et al. 2024; Plessy et al. 2024; Schultz et al. 2024; Vargas-Chávez et al. 2025).

The increased lability of siphonophore genomes relative to other hydrozoans could have a variety of causes, both neutral and adaptive. Demographic changes could lead to relaxed selection that results in genome expansion and reorganization (Lynch 2007). Often, a high amount of heterogeneous repeats in other organisms are also a result of both accumulation and slow rates of removal of repetitive DNA (Kelly et al. 2015). Alternatively, particular changes in genome architecture could be associated with specific changes in the biology of siphonophores as has been similarly proposed in clitellates with the change from marine to freshwater (Lewin et al. 2024; Schultz et al. 2024; Vargas-Chávez et al. 2025). The challenge is that siphonophores differ greatly from other animals, including their closest relatives, in a great number of ways including colonial development and a pelagic lifestyle. This makes it particularly challenging to associate specific genomic changes with specific changes in other aspects of their biology.

One of the most striking unique features of siphonophore biology is the extreme form of functional specialization between bodies (zooids) within siphonophore colonies, and the precise organization of these bodies into species-specific patterns. This unique morphology arises through unique developmental processes. Little is known, though, about the molecular mechanisms controlling these novel developmental processes. They could be completely novel to siphonophores, or the unique colony-level development of siphonophores could be coordinated with slight modifications to the same pathways that are shared with other zooid-budding cnidarians in order to be pelagic. We examined two distinct features of genome organization that could be associated with developmental novelty. First, we used phylogenetic analyses of gene annotations to see if several gene families known to be central to cnidarian development (*homeobox,* etc) have expanded in *Nanomia septata*. These gene families have not expanded and have around the same number of *homeobox* and *wnt* genes found in other closely related hydrozoans (Supplementary Table 1, Supplementary Figure 1-2). Second, we asked whether the extensive genome rearrangements found in *Nanomia septata* had brought genes that have similar expression across specialized zooids into closer physical proximity, perhaps for shared transcriptional regulation. Again, they have not (Supplementary Figure 4). The only exception are male gonophores, where we found genes relevant to spermatogenesis to be colocalized in multiple regions of the genome. This is not a siphonophore specific feature, though, and likely reflects the unique transcriptional demands of sperm as their genomes are packed (Wang et al. 2001; Boutanaev et al. 2002; Roy et al. 2002; Miller et al. 2004). This leaves it unclear what, if any, relationship there is between the radically changed genomes of siphonophores and their radically unique morphology and development. Mechanistic developmental work in siphonophores will be critical to bridging this gap in understanding.

Great strides have been made in siphonophore systematics in recent years, including a recent study of *Nanomia* (Hosia et al. 2024). That work was based on morphology and phylogenetic analyses of two genes, and newly described the species our reference genome animal belongs to as *Nanomia septata*. Individuals of *Nanomia septata* had previously not been recognized as distinct from *Nanomia bijuga*. We sequenced 20 additional genomes of *Nanomia* and mapped them to the reference scale presented here (Figure 4). This corroborates the findings of Hosia et al. (2024), adding support and new details, including regions of overlaping distribution of *Nanomia* species. *Nanomia bijuga* is found in multiple areas in the Atlantic, Mediterranean and Pacific. *Nanomia cara* overlaps with *N.bijuga* in Rhode Island, while *Nanomia* sp. 1 overlaps with *N. bijuga* in Hawai’i. These two regions harbor sympatric *Nanomia* species as there are no obvious oceanic barriers. Though describing any specific species ranges is out of the scope of this project. These results are consistent with other recent studies (Johnson et al. 2022; Church et al. 2024) that show that the diversity and geographic structure of gelatinous zooplankton has been greatly underestimated.

Understanding the specific features of global *Nanomia* diversity also has important practical importance as *Nanomia septata* emerges as an important model species for the study of development and other aspects of siphonophore biology. One of the most difficult parts of studying siphonophores is acquiring them since they must be regularly collected from the field. This makes it critical to know the diversity within the group, including how many species there are, where and when they are found, and how to tell them apart.

The chromosome-scale assembly presented here, along with its comparison to other cnidarian genomes and use as a reference for population studies, will facilitate future work on other aspects of siphonophore biology, including their unique development and morphology.

## Methods

### Genome Description

#### Specimen collection

Supplementary Table 8 presents sample data, including collection date, location and depth, for each newly sequenced specimen considered in this study. The chromosome scale genome is from a single specimen, deposited at the Yale Peabody Museum as YPM-IZ-111756 (also known as D1399-SS10-NA-1-Nanomiabijuga), collected by ROV Doc Ricketts October 30, 202 at 447 meters depth, latitude 36.7N, longitude 121.0672W in California. It was snap frozen by liquid nitrogen.

#### Karyotype

The karyotyping protocol was adapted from (Guo et al. 2018) with modifications as described below. We dissected the nectosomal and siphosomal growth zones (NGZ and SGZ) and basigasters (the base of gastrozooids closest to the stem and the origin point of cnidocytes) from four *Nanomia sp.* (*Nanomia* V4433 and *Nanomia* specimen 1,2,3 from diffusion tube culture-no sampler number available) and placed them into small petri dishes (60mm). Several basigasters were put into 1 hour colchicine treatment in the dark and the remaining basigasters, NGZ, and SGZ were put into a 5 hour colchicine treatment in the dark. Half of the basigasters from the 1 hour colchicine treatment were washed with DI water, by adding water in via drops and immediately pipetting it up. The rest of the basigasters in the 5-hour colchicine incubation were split up and washed with either a mix of DI water: filtered sea water (1:1) ratio or only DI water. Both were immediately pipetted up. We made Carnoy’s Fix fresh and added (3:1 methanol: glacial acetic acid) enough to cover the entire petri dish, which was then put on wet ice for 30 minutes.

Samples in petri dishes were transferred to slides and the liquid was removed. 20µl of 60% glacial acetic acid were added over the tissue, which was covered with siliconized coverslips using Sigmacote. We squashed preps manually using maximum pressure. Squashed preps were then placed overnight in a humidified chamber within a 4 degree refrigerator, and sealed with parafilm. The next day, siliconized coverslips were removed, samples were placed on dry ice, and each slide was washed with ∼2mL 1X PBS to cover samples and then stained with DAPI 1:5000. Slides were imaged via confocal.

#### Transcriptome sequencing

New transcriptomes were sequenced for two specimens, fresh tissue of *Nanomia septata* (Nanomia-culture-1) and frozen tissue of *Nanomia bijuga* (YPM No. 110217). We extracted RNA with the RNAqueous Total RNA Isolation Kit (Invitrogen) following the standard protocol as described in Church et al (2024). We then removed DNA using the Turbo DNAse kit (Invitrogen).

Our samples were sent to Keck Microarray Shared Resource at Yale University for PacBio library prep and Iso-Sequencing, which is as follows. 400 ng of total RNA with a RIN of >7 was used as input. First-strand cDNA synthesis and PCR amplification of cDNA products were performed using the NEBNext Single Cell/Low Input cDNA Synthesis & Amplification Module (NEB kit), in conjunction with the Iso-Seq Express Template Switching Oligo and cDNA PCR Primer from the PacBio Iso-Seq Express Oligo Kit (101-737-500), following the PacBio Iso-Seq protocol. A minimum of 200 ng of amplified cDNA was used to prepare SMRTbell libraries with the SMRTbell Prep Kit 3.0 (102-396-000). The SMRTbell libraries were prepared according to the SMRT Link Sample Setup instructions, using the Polymerase Binding Sequel® II Binding Kit 3.1, and loaded onto the Sequel II for sequencing.

Delivered reads were put through our lab’s treeinform workflow as described in Church et al (2024) and at https://github.com/dunnlab/isoseq.

#### Genome sequencing and initial assembly

Genome HiFi sequencing, Dovetail Omni-C proximity ligation, initial assembly, and annotation were completed by Cantata Bio on specimen YPM-IZ-111756 (sampling number D1399-SS10-NA-1).

Two combined DNA preps for the *Nanomia septata* sample were made, both of which used a CTAB protocol followed by the Qiagen mini kit. For the first prep, 180 mgs of sample incubated in 10 mls of CTAB, which yielded 8860ng of DNA. For the second prep, 500 mgs of sample incubated in 20 mls of CTAB, which yielded 8440 ng of DNA.DNA samples were quantified using Qubit 2.0 Fluorometer (Life Technologies, Carlsbad, CA, USA). The PacBio SMRTbell library (∼20kb) for PacBio Sequel was constructed using SMRTbell Express Template Prep Kit 2.0 (PacBio, Menlo Park, CA, USA) using the manufacturer recommended protocol.

The library was bound to polymerase using the Sequel II Binding Kit 2.0 (PacBio) and loaded onto PacBio Sequel II. Sequencing was performed on PacBio Sequel II 8M SMRT cells (2 were used). This produced 4,568,345 HiFi reads totalling 54.2GB in length.

PacBio CCS reads were assembled with Hifiasm1 v0.15.4-r347 using default parameters. Blast results of the Hifiasm output assembly against the nt database were used as input for blobtools2 v1.1.1 (Laetsch and Blaxter 2017) and scaffolds identified as possible contamination were removed from the assembly. Finally, purge_dups3 v1.2.5 was used to remove haplotigs and contig overlaps (Guan et al. 2020).

#### Structural annotation

Chromatin was fixed in place with formaldehyde in the nucleus. Fixed chromatin was digested with DNase I and then extracted, chromatin ends were repaired and ligated to a biotinylated bridge adapter followed by proximity ligation of adapter containing ends. After proximity ligation, crosslinks were reversed and the DNA purified. Purified DNA was treated to remove biotin that was not internal to ligated fragments. Sequencing libraries were generated using NEBNext Ultra enzymes and Illumina-compatible adapters. Biotin-containing fragments were isolated using streptavidin beads before PCR enrichment of each library. The Omni-C library was sequenced on an Illumina HiSeqX platform to produce ∼30x sequence coverage.

The *de novo* HiFi assembly and Dovetail OmniC library reads were used as input data for HiRise, a software pipeline designed specifically for using proximity ligation data to scaffold genome assemblies (Cheng et al. 2021). Dovetail OmniC library sequences were aligned to the draft input assembly using bwa (https://github.com/lh3/bwa). The separations of Dovetail OmniC read pairs mapped within draft scaffolds were analyzed by HiRise to produce a likelihood model for genomic distance between read pairs, and the model was used to identify and break putative misjoins, to score prospective joins, and make joins above a threshold (Lieberman-Aiden et al. 2009).

#### Assembly refinement and Quality assessment

Juicebox (JBAT) (Durand et al. 2016) was used to manually curate and fix mis-assemblies and re-orient some contigs. We followed a similar pipeline used to manually curate the *Octopus vulgaris* genome (Destanović et al. 2023), including using Chromap (Zhang et al. 2021) to map the Omni-C data with a quality cutoff of Q0 and creating .hic files using 3d-dna (Dudchenko et al. 2017) and artisanal (https://bitbucket.org/bredeson/artisanal) for use in Juicebox. In total, we went through 8 rounds of manual refinement of the genome. Scripts are deposited in our GitHub repository.

To review the quality of our assembly, we ran BUSCO metazoa_odb10 (Manni, Berkeley, Seppey, Simão, et al. 2021; Manni, Berkeley, Seppey, and Zdobnov 2021) on our assembly and NCBI FCS.

#### Gene annotation

Our gene annotation was completed by Cantata Bio as follows. Coding sequences from *Hydra vulgaris* (PRJNA703404), *Hydractinia symbiolongicarpus* (https://research.nhgri.nih.gov/hydractinia/), *Nematostella vectensis* (PRJNA667495) and *Rhopilema esculentum* (PRJNA523480 and PRJNA512552) were used to train the initial *ab initio* model for *Nanomia septata* using the AUGUSTUS software v2.5.5. Six rounds of prediction optimisation were done with the software package provided by AUGUSTUS. The same coding sequences were also used to train a separate *ab initio* model for *Nanomia septata* using SNAP v2006-07-28. RNA-seq reads were mapped onto the genome using the STAR aligner software v2.7 and intron hints generated with the bam2hints tools within the AUGUSTUS software. MAKER, SNAP and AUGUSTUS (with intron-exon boundary hints provided from RNA-seq) were then used to predict for genes in the repeat-masked reference genome. To help guide the prediction process, Swiss-Prot peptide sequences from the UniProt database were downloaded and used in conjunction with the protein sequences from *Hydra vulgaris, Hydractinia symbiolongicarpus, Nematostella vectensis* and *Rhopilema esculentum* to generate peptide evidence in the Maker pipeline. Only genes that were predicted by both SNAP and AUGUSTUS softwares were retained in the final gene sets. To help assess the quality of the gene prediction, AED scores were generated for each of the predicted genes as part of the MAKER pipeline. Genes were further characterized for their putative function by performing a BLAST search of the peptide sequences against the UniProt database. tRNA were predicted using the software tRNAscan-SE v2.05.

### Gene trees

We developed a Snakemake workflow to build gene trees using proteomes from *N. septata*, 2 choanoflagellates, and 19 other animals (porifera, ctenophore, placozoa, ecdysozoa, spiralia, and chordata; listed in the github repo under: “analyses_annotation/gene_trees/config/download_targets.tsv”). The workflow downloads protein sequences, functionally annotates them using eggNOG-mapper v2.1.10 (Huerta-Cepas et al. 2019; Cantalapiedra et al. 2021) and infers individual gene trees using Orthofinder v2.5.4 (Emms and Kelly 2015; Emms and Kelly 2019) with default parameters. It merges all gene trees in a master text file. We identified key genes that are conserved across Metazoa, with a focus on Cnidaria, including *Homeobox* and *Wnt* genes. We identified orthogroups containing homeobox sequences using human homeobox genes functionally annotated by eggNOG-mapper as landmarks. We corroborated *N. septata* orthofinder results by identifying homeodomain-containing proteins running HMMER v3.4 (hmmer.org) with the hidden Markov model hd60.hmm from Zwarycz et al. (2015), following Steinworth et al. (2022). In order to accurately identify Wnt and Hox-related genes, we used their gene trees from Orthofinder as a starting point for a more detailed phylogenetic analysis.

We retrieved 1,114 protein sequences from the orthogroup containing Antp homeobox genes and combined them with 111 previously identified Hox/Hox-related sequences from Khalturin et al. (2019). We aligned these 1,225 sequences with the L-INS-i algorithm using MAFFT v7 (Katoh and Standley 2013), which accounts for one conserved domain and long gaps. After manually trimming the variable regions and keeping only the 60aa homeodomain, we added 270 Hox/Hox-related homeodomain sequences from Steinworth et al. (2023) using MAFFT with the option –add fragments. We trimmed this alignment using trimAl (options -gt 0.9 -seqoverlap 90 -resoverlap 0.1) (Capella-Gutiérrez et al. 2009) and inferred a maximum likelihood tree using IQTree2 (Minh et al. 2020) with the evolutionary model LG+R8, selected by ModelFinder (Kalyaanamoorthy et al. 2017). Ultrafast bootstrap branch support values were calculated based on 1,000 replicates. This tree was rooted on the stem branch of Nk-like genes.

For inferring the Wnt gene tree, we first combined all protein sequences assigned to the *Wnt* orthogroup (317 sequences), 13 *Wnt* sequences from the hydrozoan jellyfish *Clytia hemisphaerica* previously identified by Condamine et al. (2019), and 12 *Wnt* sequences from the cubozoan jellyfish *Tripedalia* identified by Khalturin et al. (2019). We removed two sequences from *Orbicella faveolata* that were exceptionally long (XP_020618804.1 and XP_020632229.1). We aligned the remaining 342 sequences and 7 DASH sequences (for structural homology, see (Rozewicki et al. 2019) using the FFT-INS-ii algorithm on MAFFT v7 (Katoh and Standley 2013). Using IQTree2 (Minh et al. 2020) (Minh et al. 2020), we selected the best evolutionary model (LG+R8) and inferred a maximum likelihood tree, with ultrafast bootstrap branch support values based on 1,000 replicates.

Additional *Homeobox* and *Wnt* genes that we found manually (not originally included in the annotation) have been added to annotation files (transcript, protein and gff). These sequences were found in the *Nanomia septata* iso-sequencing and were checked before their addition.

#### Differential expression and Zooid covariance/co-expression analyses

We re-examined previously published zooid-specific bulk RNA-seq data (Munro et al. 2022) using salmon v1.10.1 (Patro et al. 2017). Counts were then imported with tximport v1.26.1 into DESeq2 v1.38.3. Our scripts also produced figures to show the variance/comparison between different zooid types and developmental stages and how they relate to one another (Supplementary Figure 3).

We next sought to identify any genes that are close to each other physically and have strong expression covariance across zooids. This analysis was implemented in the script in the repository https://github.com/dunnlab/nanomia_genome/blob/main/analyses_de/expression-proximity.ipynb. First, for each gene we average the normalized counts across replicates for each combination of tissue and stage that was sequenced. This generated a dataframe with one row per gene and one column per tissue and stage. We ordered genes by chromosome and start site. The gene expression covariance matrix was calculated based on the averaged normalized expression values. We then created a physical distance matrix between genes, where the distance is nan if the genes are on different chromosomes and the number of genes away if they are on the same chromosome, eg 0 for self, 1 for adjacent, etc. We then derived a proximity matrix from this distance matrix by passing it through a gaussian kernel and taking the inverse. In the proximity matrix, the value is 0 if the genes are on different chromosomes, 1 on the diagonal (ie, the proximity of a gene to itself is 1), and has a smooth transition from 1 to 0 as the genes become more distant. The standard deviation of the kernel was set such that proximity is 0.5 for genes that have a distance of 50. Finally, we took the element-wise product of the expression covariance matrix and the proximity matrix, producing a matrix we refer to as the expression proximity product. Values in this expression proximity product are near zero if the genes are far apart or have low correlation. They are near one if they are physically close and have high positive correlation, and near negative one if they are physically close and have strong negative correlation.

#### Custom repetitive element library generation and TE detection

Since the *Nanomia septata* genome presented here is the first genome assembly for the genus *Nanomia*, the number of repetitive elements registered in databases, such as Dfam, is limited relative to typical model animals (Storer et al. 2021). Thus, we generated a custom repeat library using RepeatModeler v2.0.5 (Flynn et al. 2020; Storer et al. 2021). Inactivated TEs obtain the accumulation of mutations, including base substitutions, insertions, and deletions (Goubert et al. 2022; Matsushima et al. 2024). Because of this, when RepeatClassifier, a submodule of RepeatModeler, performs a search against Dfam using rmblast, the number of hits can be extremely low, resulting in a library containing many uncharacterized sequences (“Unknown” repeats). This is particularly evident for non-model species whose repetitive elements are not well-represented in Dfam. To address this challenge, we used an approach similar to that used in our previous studies (Flynn et al. 2020; Kon-Nanjo et al. 2024). First, we generated a custom repetitive element library using RepeatModeler v2.0.5 (Flynn et al. 2020) with the *Nanomia septata* genome assembly as the query. Then, we obtained all Dfam HMM profiles of TEs from the Dfam database (https://dfam.org/home). Because the Dfam HMM profile dataset for TEs is a huge data size, we divided these HMM profiles into smaller chunks, each containing 25,000 HMM profiles. Each chunk of HMM profiles was indexed using hmmpress (Neuwald and Poleksic 2000). Then, we performed an HMM-based search against the Dfam HMM profiles using nhmm scan with default parameters, using the custom repeat library as the query (Neuwald and Poleksic 2000). For each query sequence, unclassified query sequences were assigned names using the annotation of the hit with the lowest E-value (≤ 0.05). Using the updated custom repetitive element library, we conducted RepeatMasker (Hausmann and Kurtz 2021) on the *Nanomia septata* genome assembly with the options of -parallel 70 -gff -a -dir -xsmall. For other species, we obtained genome assemblies from the NCBI genome database, and performed RepeatModeler and RepeatMasker in the same manner.

### Comparative cnidarian genomics

#### Synteny

We performed synteny analysis using odp v0.3.3 (Simakov et al. 2022) on the hap2 *Nanomia septata* genome against other publicly available cnidarian chromosome-scale genomes with a proteome and available annotation (*Nematostella vectensis* GCF_932526225.1, *Rhopilema esculentum* (Schultz et al. 2023), *Hydra vulgaris* GCA_022113875.1, and *Hydractinia symbiolongicarpus* GCF_029227915.1, and *Physalia physalis* (Li et al. 2020; Church et al. 2024). The *P. physalia* genome was annotated using the *P. physalis* isoseq data (Church et al. 2024), SRR29478154), and processed with the Dunn Lab isoseq workflow (https://github.com/dunnlab/isoseq, modified from Guang, et al (2021). We selected the transcriptome with a cutoff value of 8 under strict conditions, resulting in 13048 transcripts. The transcriptome was then mapped to the genome using minimap2 v2.2 (Li 2018; Li 2021), and the resulting *bam* file was sorted using SAMtools v1.18 (Danecek et al. 2021). The *bam* file was then converted to a *gff* file using spliced_bam2gff v1.3 (Anon). The input proteome was generated by translating the mapped transcriptome using TransDecoder v5.71 (https://github.com/TransDecoder/TransDecoder). We then plotted syntenic relationships using the main odp v0.3.3 workflow between each pair of species and identified ancestral linkage groups as defined in Simakov et al. (2022) and Schultz et al (2023). We ran odp_rbh_to_ribbon on *.rbh* files containing information on bilaterian-cnidarian-sponge (BCnS) ancestral linkage groups (Schultz et al. 2023) to generate a ribbon plot.

### Population Genetics and Mitochondrial Genomes

*Nanomia* samples (sample identifiers NA28-38 and WS2-10) were sequenced via Illumina Nova seq S4 2X150 PE. Coverage ranged from 250 million reads to 900 million reads (Supplementary Figure 11). Genome size was estimated for samples NA28-38 and WS1-10, for which there was sufficient coverage to achieve an accurate estimate. For this analysis, we used a Snakemake workflow (Köster and Rahmann 2018) to count k-mers using Jellyfish v2.3.0 (Marçais and Kingsford 2011) and Genomescope v2.0 (Vurture et al. 2017; Ranallo-Benavidez et al. 2020) as detailed in Ahuja et al. (2024).

For all samples, libraries were trimmed for Illumina adapters using Trimmomatic v0.39 (Bolger et al. 2014) and library quality was assessed using FastQC v.0.11.9 (Andrews 2010). Mitochondrial genomes were assembled from a subset of 10,000,000 read pairs using GetOrganelle v.1.7.7.0 (Jin et al. 2020), using as a seed the SRR23143273 and SRR23143286 publicly available mt genomes from *Nanomia bijuga* sample CWD116 and *Nanomia* sp.

California NA19 (Ahuja et al. 2024), and using the animal_mt database and default parameters. All mitochondrial genomes were assembled as linear, with the exception of YPM IZ 35039; upon visual inspection of mitogenome alignment, this was considered to be a misassembly, and we manually corrected this genome. The mitogenome for YPM IZ 107008 was assembled in three fragments. For downstream analyses the largest fragment was used. Mitochondrial genomes were annotated with MITOS2 (Donath et al. 2019) and tRNAscan (Chan et al. 2021).

A mitochondrial genome was also assembled from PacBIo HiFi reads for the genome reference specimen by blasting CO1 from the 2 seed *Nanomia* mt genomes and identifying the appropriate contigs. Assembled mitochondrial sequences were combined with publicly available mt genomes for *Nanomia* (OQ957219.1, OQ957196.1), *Halistemma* (OQ957197.1, OQ957195.1), and *Agalma* (OQ957203.1, OQ957191.1, OQ957190.1, OQ957189.1,), and then aligned using MAFFT, --adjustdirectionaccurately option v.7.505 (Katoh and Standley 2013) and a phylogeny was inferred using IQTree2 software (Minh et al. 2020), model autoselected (Kalyaanamoorthy et al. 2017) and 1,000 ultrafast bootstraps (Hoang et al. 2018). Assembled mitochondrial sequences are available on our GitHub repository (https://github.com/dunnlab/nanomia_genome).

In addition, individual marker sequences for the mitochondrial cytochrome oxidase I (CO1) gene were assembled from a subset of 10 million reads using *in silico* PCR as implemented in sharkmer (available at https://github.com/caseywdunn/sharkmer, commit ‘c43cfc2’). Sequences were combined with publicly available *Nanomia* sequences, as published in Hosia et al (2024), along with sequences for *Halistemma* and *Agalma* from NCBI. Sequences were aligned with MAFFT, and gene trees inferred with IQTree2, as described above.

Assembled CO1 sequences are available on our GitHub repository.

Population structure was analyzed following the workflow described in Church et al (2024). In brief: libraries were assessed for contamination with Kraken2 (Wood and Salzberg 2014); sample kinship was evaluated using Plink2 (Chang et al. 2015); reads were then mapped to the reference assemble using BWA (Li and Durbin 2009) and alleles were called using BCFTools v1.16 (Li 2011); genotype likelihoods were calculated using ANGSD v.0.935 (Korneliussen et al. 2014); principal components of genomic variation were analyzed on genotype likelihoods using PCANGSD v1.21 (Skotte et al. 2013). We repeated this analysis for *N. bijuga* and *N. septata* separately, using assembled transcriptomes from PacBio Iso-Seq data as the reference, and compared our results to those using the *N. septata* reference genome.

### Data availability

Scripts and code are available at https://github.com/dunnlab/nanomia_genome. Raw reads for *Nanomia septata* genome assembly and annotation were deposited to NCBI SRA as BioProject PRJNA971210. *Nanomia septata* genome assemblies are under accession numbers: JAVHZB000000000 and JAVHZC000000000. The haplotype 1 (alternate) assembly is under BioProject PRJNA990459. The haplotype 2 (primary) assembly is under BioProject PRJNA990460. *Nanomia septata* annotation files and *Nanomia* mitochondrial genomes are also in the github repository. Illumina reads from *Nanomia* specimens have been deposited on NCBI SRA under BioProjects PRJNA1252167 (NA28-38 and WS2-10) and PRJNA925656 (NA19).

## Acknowledgements

We thank the Yale Center for Genome Analysis and Keck Microarray Shared Resource at Yale University for providing the necessary PacBio sequencing services and Illumina sequencing services, which is funded in part by the National Institutes of Health instrument grant 1S10OD028669-01 and 1S10OD030363-01A. Thank you to NIH T32 Training Grant in Genetics. We also had support from the National Science Foundation grant NSF-OCE 1829835, and the Tal Waterman Fund. This work was also funded by the Austrian Science Fund FWF (I4353) and ERC-H2020-EURIP grant number 945026 to OS, Takeda Science Foundation to TK, the Mochida Memorial Foundation for Medical and Pharmaceutical Research to TK, the International Medical Research Foundation to TK, the Yamada Science Foundation to TK. Sample collections were supported by the David and Lucile Packard Foundation. Phylopic of *P.physalis* is from: https://www.phylopic.org/images/6872620a-34e1-4744-b536-a5b5437b9097/physalia-physalis Illustration by Noah Schlottman (2013), photo by Casey Dunn (2005). Illustration under CC BY-SA 3.0 license. No further modifications were made.

## Author Contributions

NA-assembly, karyotyping, manuscript writing, differential gene expression analysis, sample collection, DNA extractions

DS- assembly, synteny

DD- assembly, synteny analysis

SHC- pop gen, mito genomes

NP- annotation, gene trees

CM- zooid differential analysis, help with karyotyping protocol/adjustments

CWD- project conception, assembly, expression analyses, manuscript writing

WS- DNA extractions

SH- sample collection

MM- karyotyping

KK- repeat analysis annotation

TK- repeat analysis annotation

OS- repeat analysis annotation

## Supplement

Supplementary Table 1. Number of sequences placed in orthogroups containing *homeobox* genes.

Supplementary Table 2. Differentially expressed genes for all zooid types and stages in *Nanomia septata*

Supplementary Table 3. Differential expressed genes that are shared between some zooid types in *Nanomia septata*.

Supplementary Table 4. Genome coverage of major TE groups Supplementary Table 5. Genome coverage of TE families

Supplementary Table 6: Support values for synteny analyses for Figure 3 and Supplementary Figure 7

Supplementary Table 7: The names used for chromosomes in the manuscript vs. their accession number/sequence name

Supplementary Table 8. Specimen data and accession numbers of all WGS *Nanomia* samples that are introduced in this study.

Supplementary Figure 1. Antp homeobox gene tree (Hox and Nk-related gene families).

Supplementary Figure 2. Gene Trees for *Wnt* genes

Supplementary Figure 3. Differentially expressed gene analysis for *Nanomia septata*.

Supplementary Figure 4. Expression proximity product for genes on each *Nanomia septata* chromosome.

Supplementary Figure 5. TE expansions in the *Nanomia septata* genome.

Supplementary Figure 6. Genome coverages of TEs.

Supplementary Figure 7. BCnS ALGs and unicellular-metazoan (pre-metazoan) ALGs in *Hydractinia symbiolongicarpus, Nanomia septata and Physalia physalis*.

Supplementary Figure 8. CO1 *Nanomia* tree.

Supplementary Figure 9. Genomescope fit models for 1 representative *Nanomia* sample per group in the phylogeny.

Supplementary Figure 10. *Nanomia* specimens mapped using 2 separate transcriptome references to look at variance.

Supplementary Figure 11. *Nanomia* sample coverage statistics.

## Notes

https://github.com/dunnlab/nanomia_genome

### Competing Interest Statement

The authors have declared no competing interest.

