## Supplementary Material for "A Chromosome-Scale Genome of *Nanomia septata* Reveals Extensive Rearrangement But No Clear Driver of the Unique Colony-Level Organization of Siphonophores"

*Nanomia* Genome Supplemental Material

Figures

Supplementary Figure 1. Antp homeobox gene tree (Hox and Nk-related gene families). Maximum likelihood tree of 1,465 Hox and Nk-related sequences, built under model LG+R8, with ultrafast bootstrap branch support (1,000 replicates) values along branches. This tree contains protein sequences from the orthogroup with Antp homeobox genes, in addition to previously identified Hox/Hox- related sequences from Khalturin *et al*. (2019) and Steinworth *et al*. (2022). *Nanomia septata* sequences are colored in magenta and highlighted with a star. Sequence names retrieved from Antp orthogroup contain the species name (followed by _pep), the sequence ID from original source file and a label with the functional annotation generated by eggNOGG-mapper. Branch lengths were transformed to equal for display. Original tree is available in the supplementary data folder.

*see separate pdf file

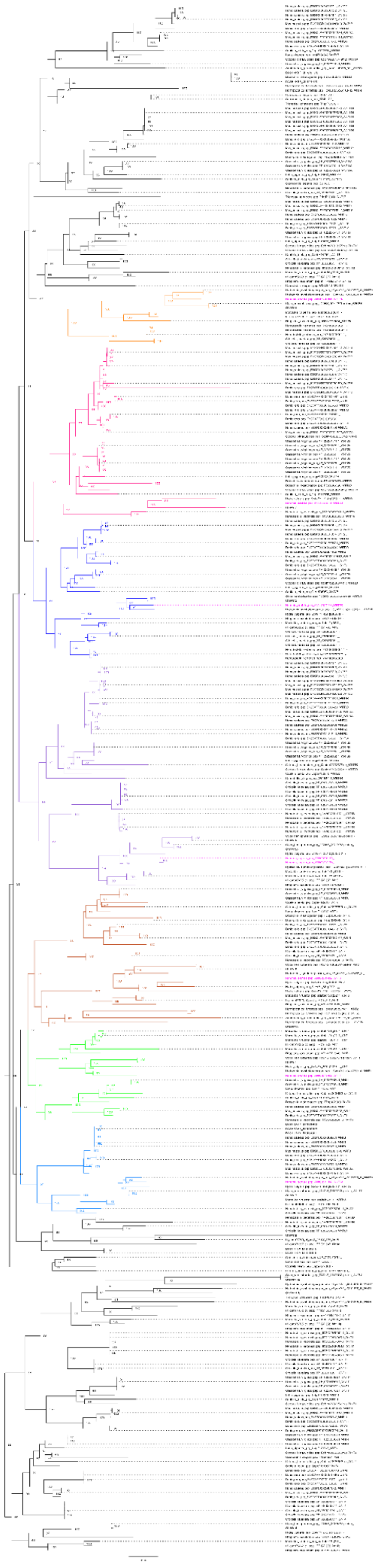

Supplementary Figure 2. *Wnt* gene tree. Maximum likelihood phylogenetic tree, midpoint rooted, inferred using the LG+R8 substitution model from 342 protein sequences gathered from representatives of Cnidaria and Bilateria. *Nanomia septata* sequences are colored in magenta. Sequence names contain the species name (followed by _pep), the sequence ID from original source file and a label with the functional annotation generated by eggNOGG-mapper. Condamine (2019) Wnt sequences from *Clytia* are labeled CheWnt1-11. Wnt sequences from *Tripedalia* previously identified by Khalturin et al. (2019) start with "'tri_comp". Ultrafast bootstrap values are displayed by their corresponding nodes.

*see separate pdf file

A

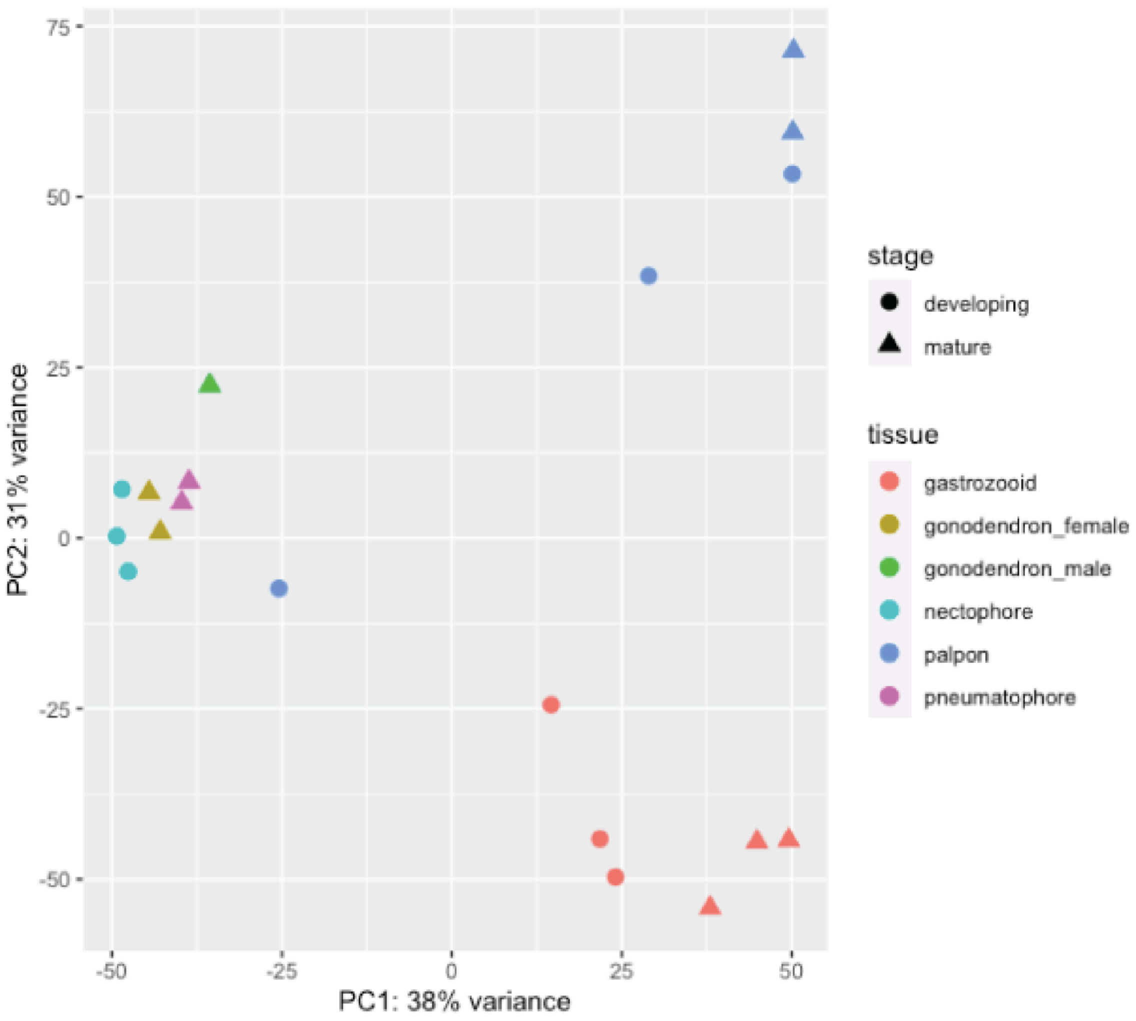


Supplementary Figure 3. Differentially expressed gene analysis for *Nanomia septata.* Transcriptomes from distinct zooid types and pneumatophores of *Nanomia septata* at different developmental stages cluster according to their transcriptional similarity.

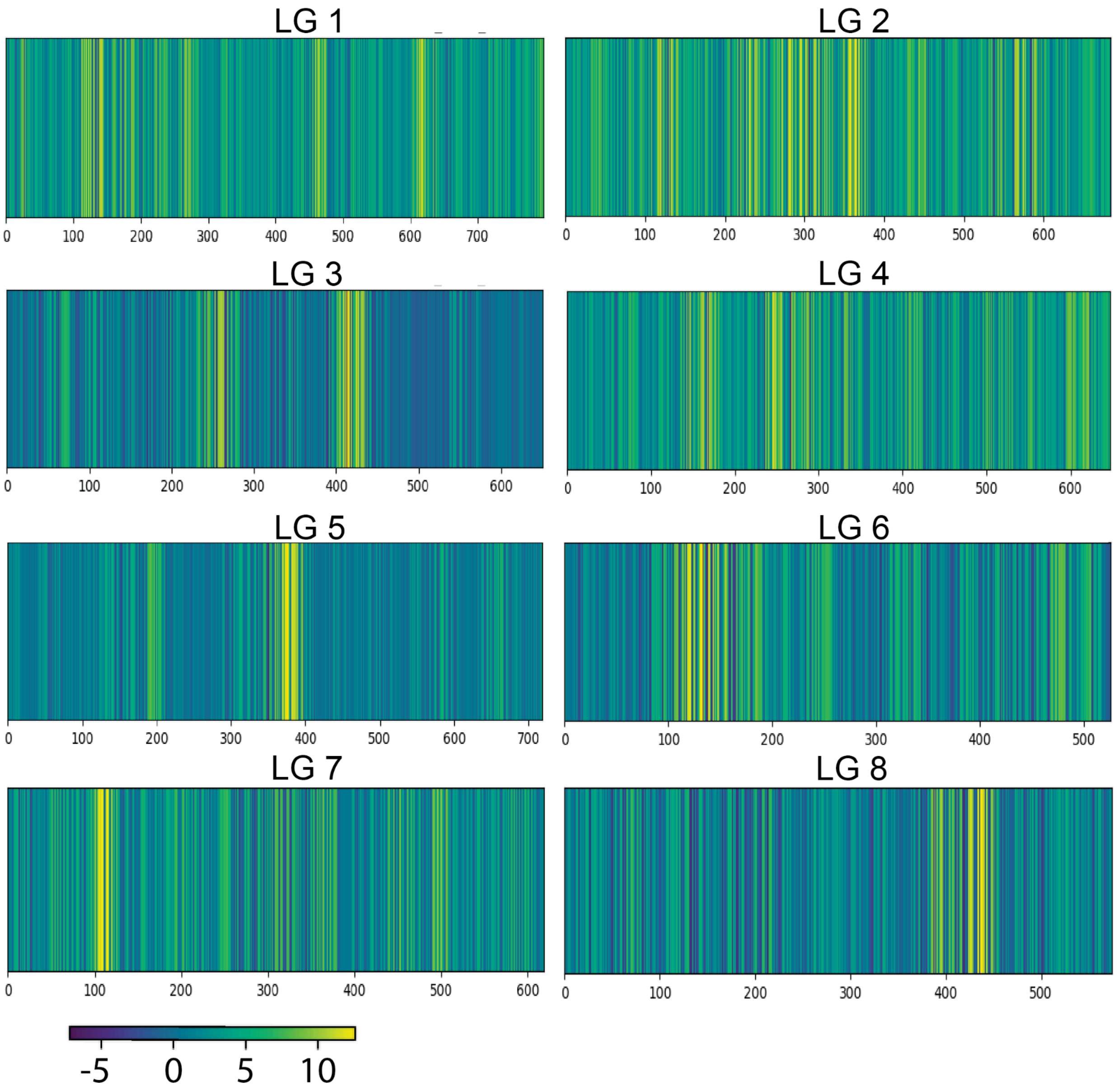


Supplementary Figure 4. Expression proximity product for genes on each *Nanomia septata* chromosome. Each panel presents a chromosome. Each bar is a gene, and the x axis indicates the index of the gene. Color indicates how strongly the expression of a gene covaries with its neighbors  (i.e. the more yellow a gene is, the more similar its expression is across zooids to neighboring genes ).

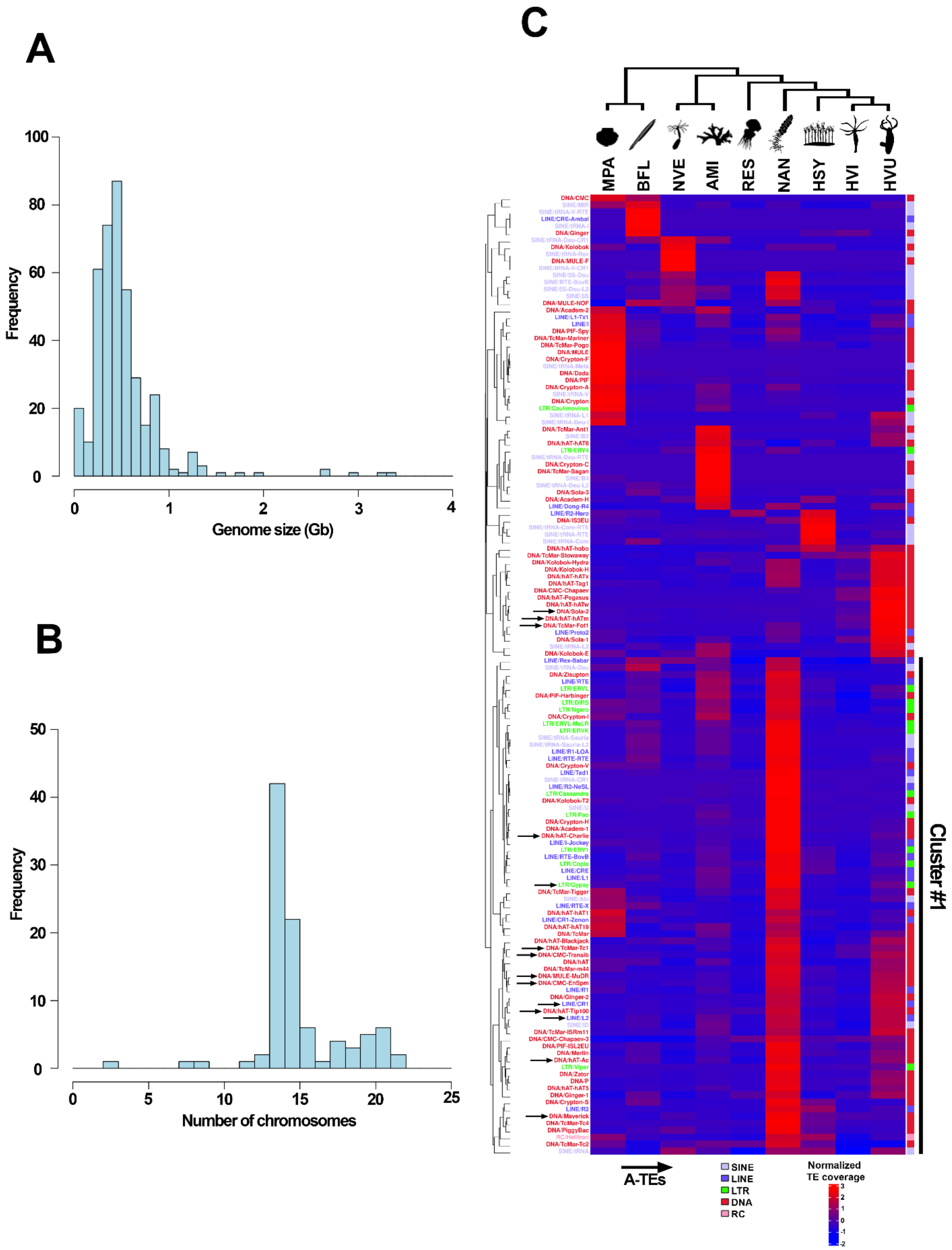
A
100
80
60
40
20
0
Frequency
0
1
2
3
4
Genome size (Gb)
B
50
40
30
20
10
0
Frequency
0
5
10
15
20
25
Number of chromosomes
C
MPA
BFL
NVE
AMI
RES
NAN
HSY
HVI
HVU
Cluster #1
A-TEs
SINE
LINE
LTR
DNA
RC
Normalized
TE coverage

Supplementary Figure 5. Transposable Element (TE) expansions in the *Nanomia septata* genome. A. Size distribution of the cnidarian genome assemblies available in the NCBI Genome database. B. Number of chromosomes in the genome assemblies available in the NCBI Genome database. C. Relative genome coverage of TE families in cnidarian genomes and two bilaterian outgroup species. For each TE family, genome coverage in each species was calculated and was normalized (z-transformed) by row (across species). The dendrogram at the left of the heatmap represents the result of hierarchical clustering performed using the Euclidean distance metric and the complete linkage method. The cluster #1 is composed of TE families which are overrepresented in nanomia compared to other species. Black arrows indicate 14 TE families belonging to the A-TEs[2]. Note that in addition to the TE families increasing in *Nanomia septata,* there are TE families showing clade-specific expansion in other animals as well. Abbreviations: PMA, *Pecten maximus*; BFL, *Branchiostoma floridae*; NVE, *Nematostella vectensis*; AMI, *Acropora millepora*; RES, *Rhopilema esculentum*; NAN, *Nanomia septata*; HSY, *Hydractinia symbiolongicarpus*; HVI, *Hydra viridissima*; HVU, *Hydra vulgaris*.

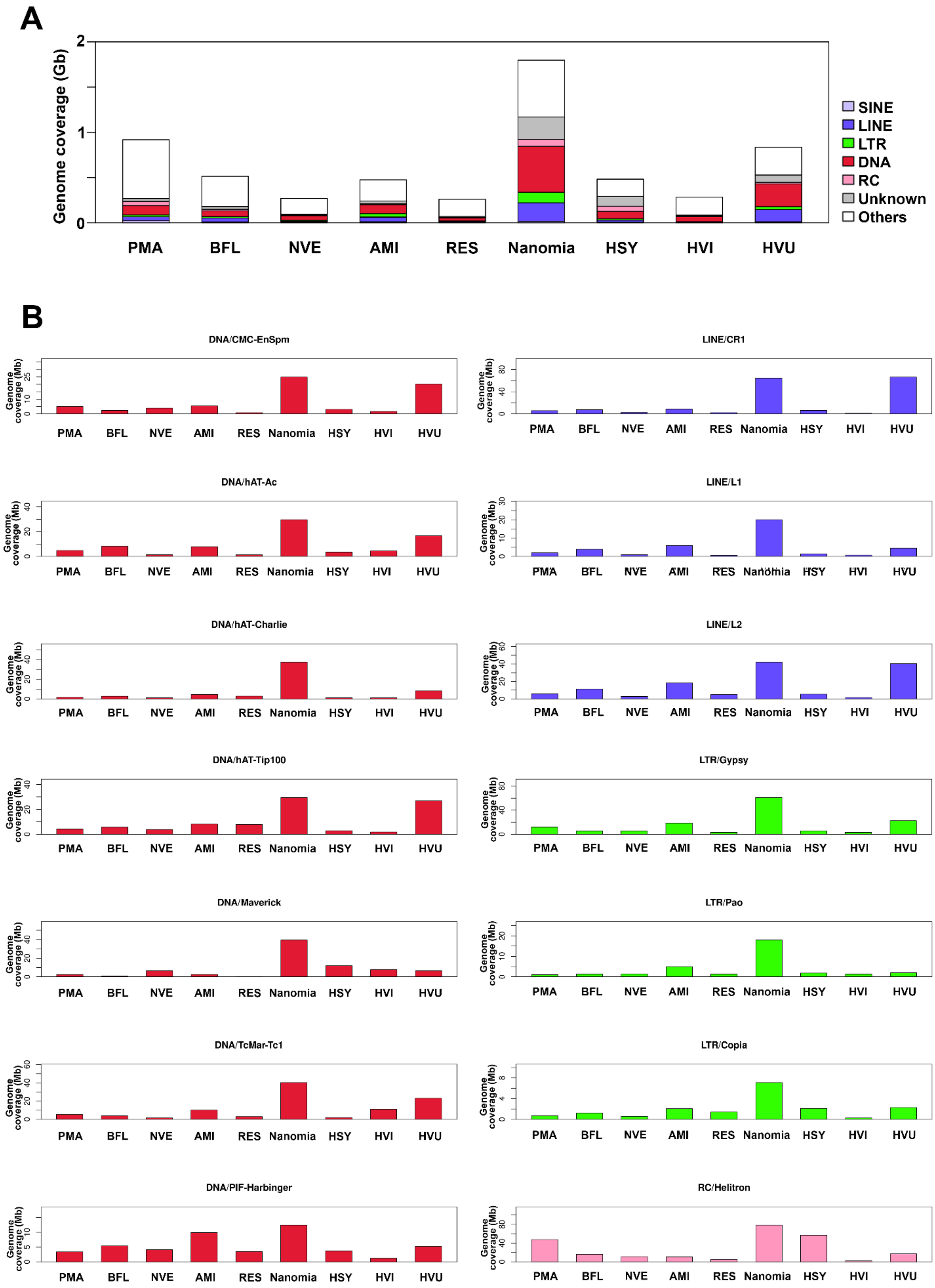
A
Genome coverage (Gb)
0
1
2
SINE
LINE
LTR
DNA
RC
Unknown
Others
PMA BFL NVE AMI RES Nanomia HSY HVI HVU
B
DNA/CMC-EnSpm
DNA/hAT-Ac
DNA/hAT-Charlie
DNA/hAT-Tip100
DNA/Maverick
DNA/TcMar-Tc1
DNA/PIF-Harbinger
LINE/CR1
LINE/L1
LINE/L2
LTR/Gypsy
LTR/Pao
LTR/Copia
RC/Helitron
Genome coverage (Mb)
PMA BFL NVE AMI RES Nanomia HSY HVI HVU

Supplementary Figure 6. Genome coverages of TEs. A. Genome coverages of SINE, LINE, LTR, DNA, and Rolling-circle elements. The y-axis indicates genome coverage of TEs (Gb). B. Genome coverages of the eukaryotic core TEs. The y-axis indicates genome coverage of TEs (Mb) and x-axis are species. The color scheme is the same as in panel a. Abbreviations: PMA, *Pecten maximus*; BFL, *Branchiostoma floridae*; NVE, *Nematostella vectensis*; AMI, *Acropora millepora*; RES, *Rhopilema esculentum*; NAN, *Nanomia septata*; HSY, *Hydractinia symbiolongicarpus*; HVI, *Hydra viridissima*; HVU, *Hydra vulgaris*.

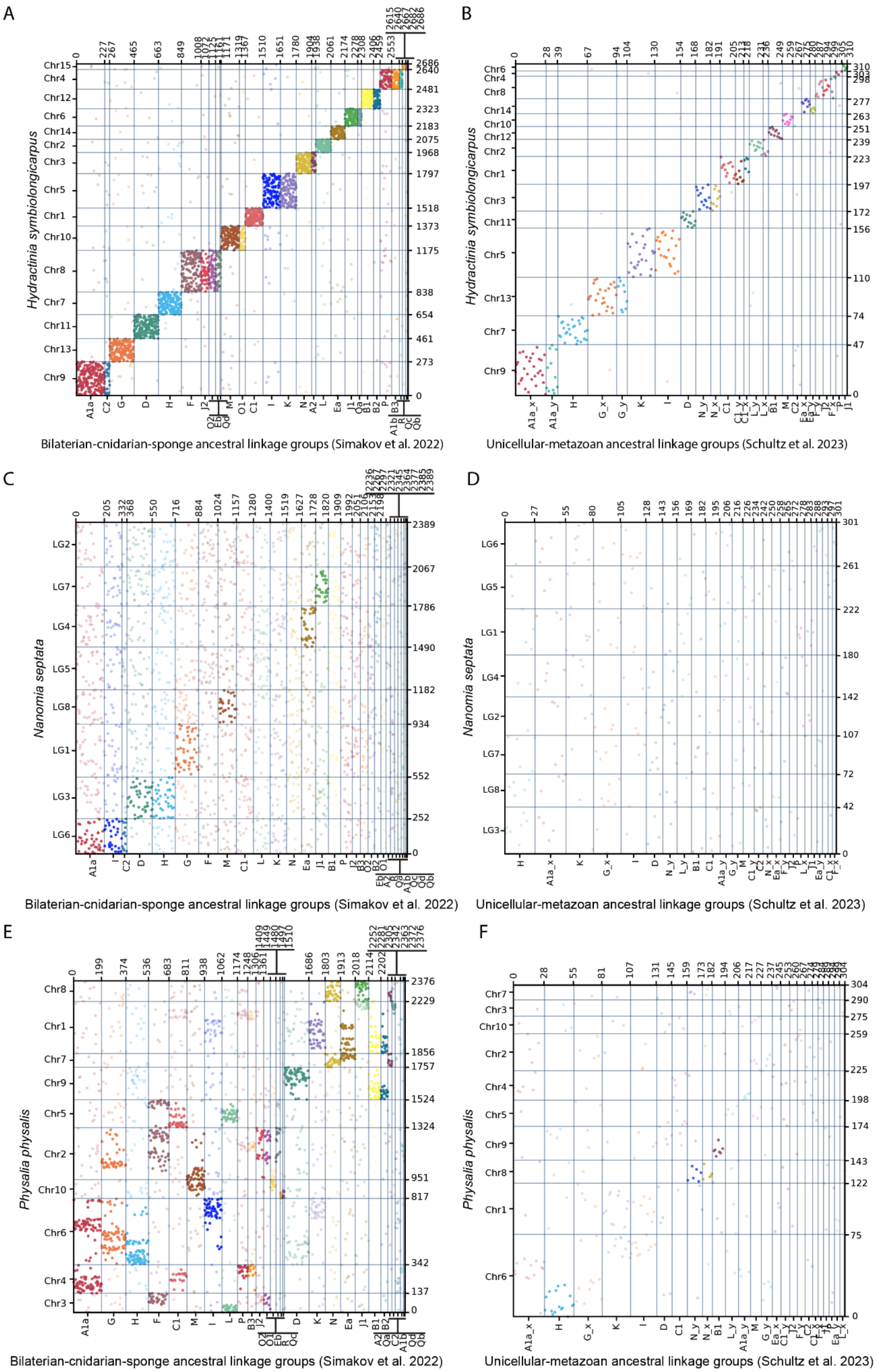
A
B
C
D
E
F
Hydractinia symbiolongicarpus
Chr15
Chr4
Chr12
Chr6
Chr14
Chr2
Chr3
Chr5
Chr1
Chr10
Chr8
Chr7
Chr11
Chr13
Chr9
Chr6
Chr4
Chr8
Chr14
Chr10
Chr12
Chr2
Chr1
Chr3
Chr11
Chr5
Chr13
Chr7
Chr9
Nanomia septata
LG2
LG7
LG4
LG5
LG8
LG1
LG3
LG6
LG6
LG5
LG1
LG4
LG2
LG7
LG8
LG3
Physalia physalis
Chr8
Chr1
Chr7
Chr9
Chr5
Chr2
Chr10
Chr6
Chr4
Chr3
Chr7
Chr3
Chr10
Chr2
Chr4
Chr5
Chr9
Chr8
Chr1
Chr6
Bilaterian-cnidarian-sponge ancestral linkage groups (Simakov et al. 2022)
Unicellular-metazoan ancestral linkage groups (Schultz et al. 2023)
Bilaterian-cnidarian-sponge ancestral linkage groups (Simakov et al. 2022)
Unicellular-metazoan ancestral linkage groups (Schultz et al. 2023)
Bilaterian-cnidarian-sponge ancestral linkage groups (Simakov et al. 2022)
Unicellular-metazoan ancestral linkage groups (Schultz et al. 2023)

Supplemental Figure 7. BCnS ALGs and unicellular-metazoan (pre-metazoan) ALGs in *Hydractinia symbiolongicarpus, Nanomia septata and Physalia physalis* shown via Oxford dot plots (ODP). A. ODP of ancestral linkage groups from Simakov et al. (2022) vs. *Hydractinia symbiolongicarpus* chromosomes. B. ODP of ancestral linkage groups from Schultz et al. (2023) vs. *Hydractinia symbiolongicarpus* chromosomes. C. ODP of ancestral linkage groups from Simakov et al. (2022) vs. *Nanomia septata* chromosomes. D. ODP of ancestral linkage groups from Schultz et al (2023) vs. *Nanomia septata* chromosomes. E. ODP of ancestral linkage groups from Simakov et al. (2022) vs. *Physalia physalis.* F. ODP of ancestral linkage groups from Schultz et al (2023) vs. *Physalia physalis.*

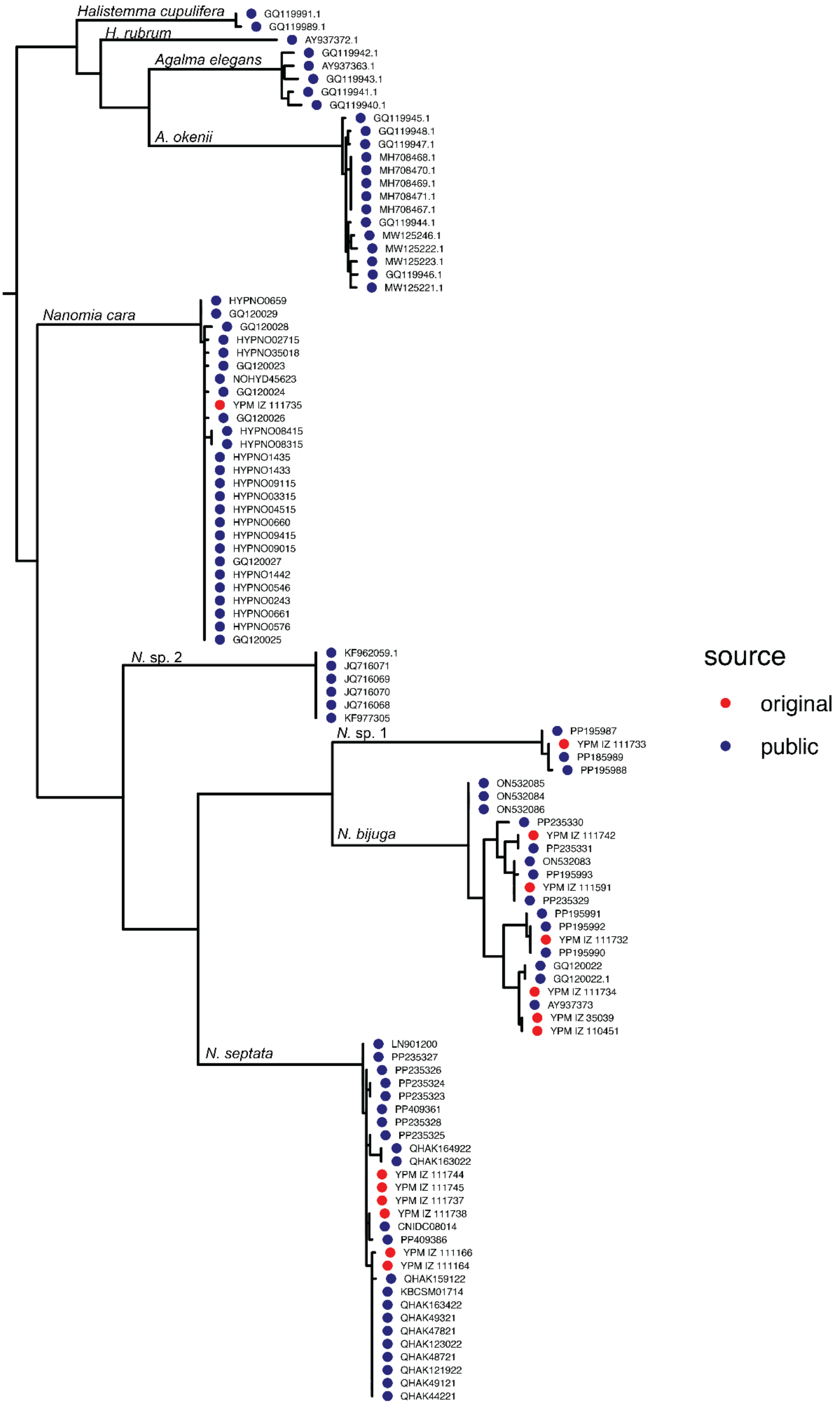
Halistemma cupulifera
GQ119991.1
GQ119989.1
H. rubrum
AY937372.1
Agalma elegans
GQ119942.1
AY937363.1
GQ119943.1
GQ119941.1
GQ119940.1
A. okenii
GQ119945.1
GQ119948.1
GQ119947.1
MH708468.1
MH708470.1
MH708469.1
MH708471.1
MH708467.1
GQ119944.1
MW125246.1
MW125222.1
MW125223.1
GQ119946.1
MW125221.1
Nanomia cara
HYPNO0659
GQ120029
GQ120028
HYPNO02715
HYPNO35018
GQ120023
NOHYD45623
GQ120024
YPM IZ 111735
GQ120026
HYPNO08415
HYPNO08315
HYPNO1435
HYPNO1433
HYPNO09115
HYPNO03315
HYPNO04515
HYPNO0660
HYPNO09415
HYPNO09015
GQ120027
HYPNO1442
HYPNO0546
HYPNO0243
HYPNO0661
HYPNO0576
GQ120025
N. sp. 2
KF962059.1
JQ716071
JQ716069
JQ716070
JQ716068
KF977305
N. sp. 1
PP195987
YPM IZ 111733
PP185989
PP195988
N. bijuga
ON532085
ON532084
ON532086
PP235330
YPM IZ 111742
PP235331
ON532083
PP195993
YPM IZ 111591
PP235329
PP195991
PP195992
YPM IZ 111732
PP195990
GQ120022
GQ120022.1
YPM IZ 111734
AY937373
YPM IZ 35039
YPM IZ 110451
N. septata
LN901200
PP235327
PP235326
PP235324
PP235323
PP409361
PP235328
PP235325
QHAK164922
QHAK163022
YPM IZ 111744
YPM IZ 111745
YPM IZ 111737
YPM IZ 111738
CNIDC08014
PP409386
YPM IZ 111166
YPM IZ 111164
QHAK159122
KBCSM01714
QHAK163422
QHAK49321
QHAK47821
QHAK123022
QHAK48721
QHAK121922
QHAK49121
QHAK44221
source
original
public

Supplementary Figure 8. CO1 *Nanomia* phylogram. Includes CO1 sequences from *Nanomia* genomes presented in the paper as well as CO1 sequences from NCBI.

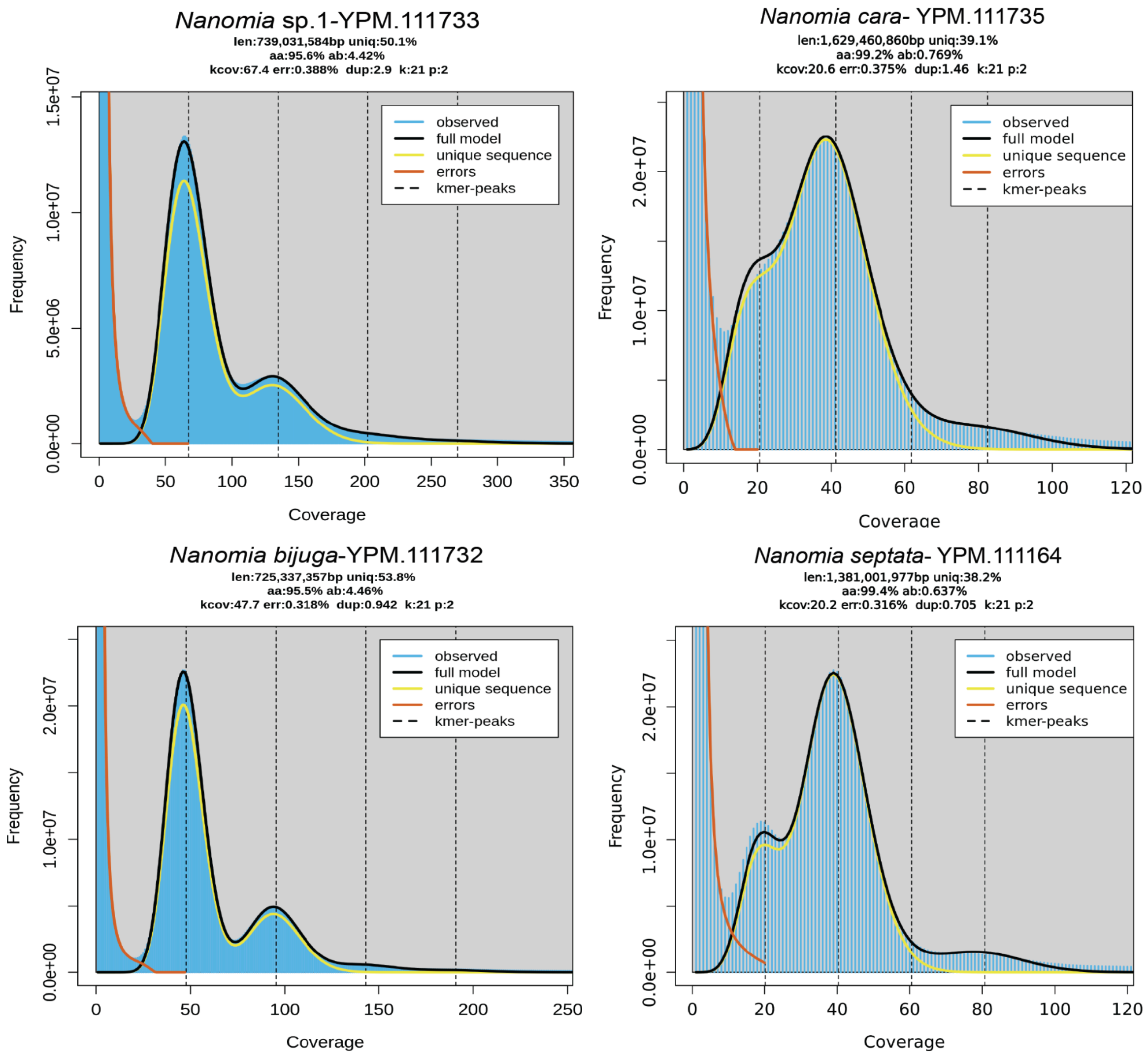


Supplementary Figure 9. Genomescope fit models for 1 representative *Nanomia* sample per group in the phylogeny, including *Nanomia septata, Nanomia bijuga, Nanomia cara* and *Nanomia* sp.1.

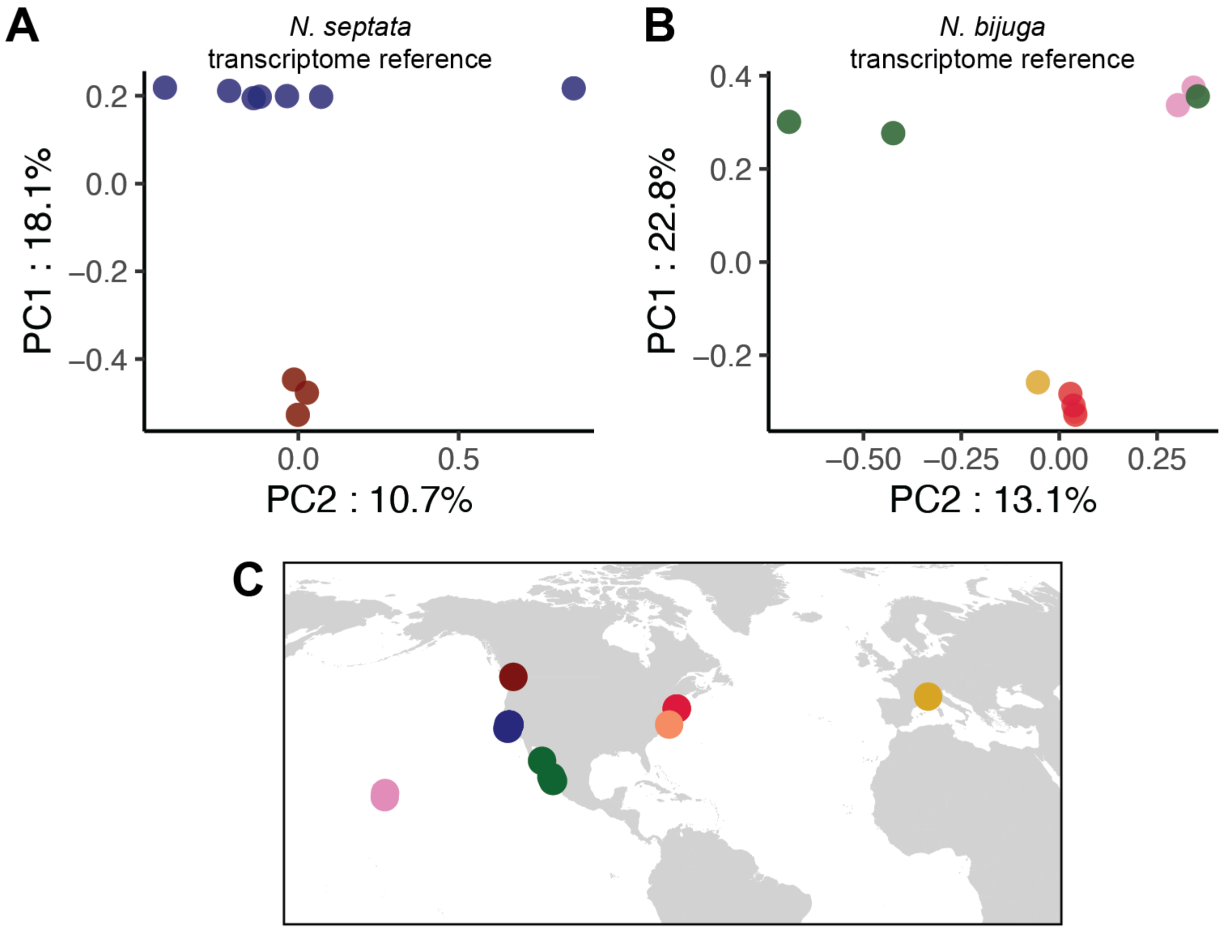


Supplementary Figure 10. *Nanomia* specimens mapped using 2 separate transcriptome references to look at variance. A. *Nanomia septata* samples mapped against *Nanomia septata* iso-seq transcriptome reference, displaying 2 distinct clusters, 1 cluster from Washington and the other from California. B. *Nanomia bijuga* samples mapped against *Nanomia bijuga* iso-seq transcriptome reference showing samples falling into multiple distinct clusters, including a cluster of Atlantic specimens + Mediterranean, a cluster of Hawaiian+GoC and 2 GoC samples that are more varied than the others. C. Map of all *Nanomia* specimen collection spots colored by population.

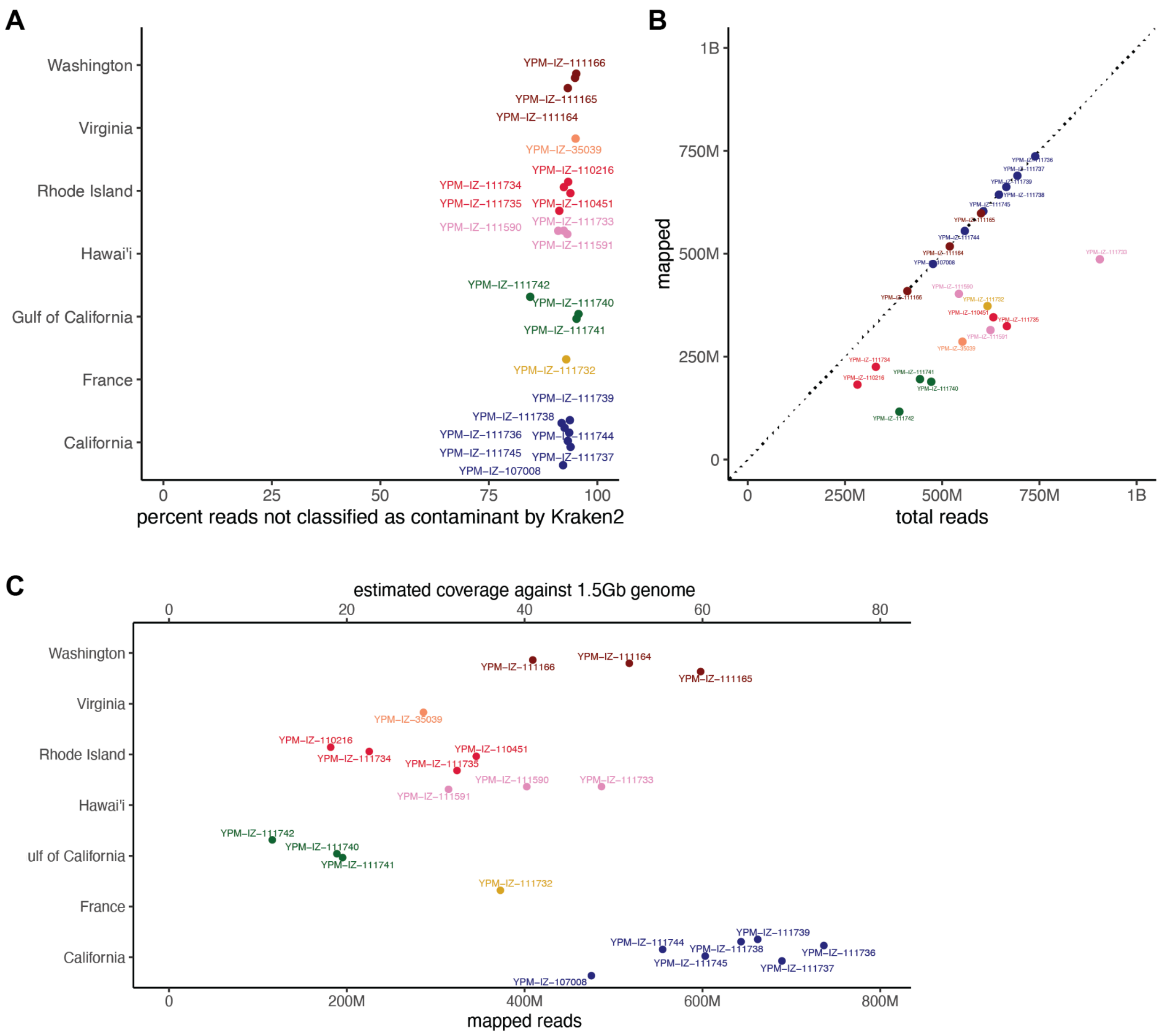


Supplementary Figure 11. *Nanomia* sample coverage statistics. A. the percent reads of *Nanomia* samples that are not classified as contamination, all falling >80%. B. The total reads for each *Nanomia* specimen. C. The total number of mapped reads against the *Nanomia septata* genome.

Tables

Supplementary Table 1. Number of sequences placed in orthogroups containing homeobox genes. These numbers refer to homeodomain containing proteins placed in orthogroups identified by searching for landmark human homeobox sequences functionally annotated by EggNOG-mapper (Cantalapiedra et al. 2021; Huerta-Cepas et al. 2019). We corroborated *N. septata* orthofinder results by identifying homeodomain-containing proteins with HMMER (hmmer.org), confirming they were placed in verified orthogroups and searching for highly divergent homeodomain-containing proteins that could have been placed in separate orthogroups. These numbers do not include sequences that were unassigned to an orthogroup or not annotated in the genome of these species. Classification of genes according to Holland et al. (2007). Orthogroup gene trees sorted by homeobox class available in the supplementary data folder. *N. septata*: *Nanomia septata*, *A. queenslandica*: *Amphimedon queenslandica*, *N. vectensis*: *Nematostella vectensis*, *M. virulenta*: *Morbakka virulenta*, *R. esculentum*: *Rhopilema esculentum*, *H. vulgaris*: *Hydra vulgaris*, *C. hemisphaerica*: *Clytia hemisphaerica*, *H. symbiolongicarpus*: *Hydractinia symbiolongicarpus*.

| Gene class | *N. septata* | *H. symbiolongicarpus* | *H. vulgaris* | *C. hemisphaerica* | *M. virulenta* | *R. esculentum* | *N. vectensis* | *A. queenslandica* |
|---|---|---|---|---|---|---|---|---|
| ANTP PRD | 33 | 42 | 40 | 46 | 69 | 53 | 113 | 14 |
| TALE | 5 | 4 | 7 | 5 | 9 | 6 | 24 | 6 |
| POU | 2 | 11 | 2 | 11 | 3 | 7 | 6 | 1 |
| LIM | 10 | 12 | 5 | 7 | 10 | 5 | 35 | 2 |
| SINE | 4 | 3 | 2 | 4 | 5 | 4 | 4 | 1 |
| CUT | 1 | 1 | 2 | 1 | 1 | 1 | 4 | 1 |
| CERS | 5 | 4 | 5 | 4 | 4 | 3 | 3 | 3 |
| HNF | 0 | 0 | 0 | 0 | 1 | 0 | 1 | 1 |
| **Total** | **60** | **77** | **63** | **78** | **102** | **79** | **190** | **29** |

Supplementary Table 2. Differentially expressed genes for all zooid types and stages in *Nanomia septata.*
*See separate tsv file

Supplementary Table 3. Differential expressed genes that are shared between some zooid types in *Nanomia septata*
*See separate tsv file

Supplementary Table 4. Genome coverage of major TE groups

| | PMA | BFL | NVE | AMI | RES | Nanomia | HSY | HVI | HVU |
|---|---|---|---|---|---|---|---|---|---|
| SINE | 24,170,260 | 10,286,797 | 6,517,676 | 11,116,136 | 1,314,613 | 15,331,805 | 6,925,188 | 305,204 | 11,023,552 |
| LINE | 42,340,798 | 42,859,608 | 12,130,717 | 52,160,374 | 11,310,953 | 205,762,304 | 22,829,424 | 5,507,125 | 135,922,726 |
| LTR | 20,407,718 | 16,006,455 | 10,327,992 | 38,327,994 | 7,143,689 | 116,102,587 | 14,387,277 | 5,720,571 | 31,202,863 |
| DNA | 102,314,463 | 63,168,007 | 50,356,689 | 98,872,614 | 34,635,363 | 507,440,070 | 82,400,611 | 56,941,786 | 251,496,006 |
| RC | 47,704,980 | 16,297,602 | 11,177,532 | 10,547,573 | 5,130,848 | 78,499,598 | 57,067,468 | 2,297,035 | 17,144,829 |
| Unknown | 32,017,292 | 30,915,358 | 3,941,283 | 29,879,779 | 12,796,988 | 246,876,936 | 108,238,438 | 13,255,870 | 81,465,677 |
| Others | 649,350,867 | 333,927,104 | 174,966,549 | 234,476,783 | 188,100,911 | 626,520,431 | 191,079,917 | 200,237,714 | 305,959,543 |

Supplementary Table 5**.** Genome coverage of TE families

| | PMA | BFL | NVE | AMI | RES | NAN | HSY | HVI | HVU |
|---|---|---|---|---|---|---|---|---|---|
| DNA/Academ-1 | 936,274 | 1,009,022 | 751,343 | 2,231,170 | 1,180,258 | 34,838,300 | 1,061,971 | 291,308 | 3,529,645 |
| DNA/Academ-2 | 2,503,621 | 54,036 | 857,499 | 2,291,450 | 261,159 | 0 | 656,736 | 0 | 76,427 |
| DNA/Academ-H | 13,083 | 0 | 60,533 | 294,199 | 0 | 0 | 142,294 | 0 | 0 |
| DNA/CMC | 102,552 | 67,272 | 0 | 0 | 0 | 0 | 0 | 0 | 0 |
| DNA/CMC-Chapaev | 89,482 | 180,275 | 41,215 | 94,506 | 13,793 | 1,081,155 | 405,970 | 2,630,275 | 8,331,676 |
| DNA/CMC-Chapaev-3 | 91,837 | 48,689 | 37,214 | 1,200,109 | 1,771,150 | 3,693,288 | 1,155,976 | 1,431,605 | 2,233,419 |
| DNA/CMC-EnSpm | 4,977,557 | 2,366,682 | 3,879,622 | 5,338,764 | 797,820 | 24,839,995 | 3,019,439 | 1,657,202 | 20,271,358 |
| DNA/CMC-Transib | 1,073,433 | 1,527,368 | 715,490 | 775,444 | 0 | 9,672,303 | 285,056 | 2,340,309 | 7,287,805 |
| DNA/Crypton | 6,460,419 | 144,665 | 0 | 1,213,147 | 0 | 463,383 | 144,978 | 24,018 | 0 |
| DNA/Crypton-A | 6,481,869 | 50,181 | 259,991 | 2,181,781 | 88,198 | 2,329,992 | 149,414 | 11,129 | 49,277 |
| DNA/Crypton-C | 0 | 0 | 0 | 117,715 | 0 | 0 | 0 | 0 | 0 |
| DNA/Crypton-F | 423,671 | 0 | 0 | 0 | 0 | 0 | 0 | 0 | 0 |
| DNA/Crypton-H | 0 | 0 | 0 | 261,004 | 110,006 | 2,003,173 | 0 | 0 | 178,314 |
| DNA/Crypton-I | 979,859 | 165,447 | 62,295 | 1,820,761 | 0 | 2,255,923 | 72,782 | 0 | 34,644 |
| DNA/Crypton-S | 0 | 72,303 | 0 | 0 | 0 | 232,078 | 75,419 | 0 | 0 |
| DNA/Crypton-V | 382,857 | 404,931 | 123,236 | 310,919 | 18,738 | 1,383,063 | 94,990 | 136,055 | 183,009 |
| DNA/Dada | 2,316,646 | 169,237 | 109,028 | 45,998 | 0 | 190,807 | 96,424 | 0 | 0 |
| DNA/Ginger | 0 | 239,152 | 0 | 0 | 0 | 0 | 37,147 | 58,382 | 0 |
| DNA/Ginger-1 | 0 | 90,094 | 0 | 30,379 | 78,714 | 241,182 | 0 | 58,382 | 176,155 |
| DNA/Ginger-2 | 0 | 7,306 | 0 | 35,836 | 8,756 | 543,001 | 37,147 | 0 | 560,721 |
| DNA/IS3EU | 3,422,094 | 1,591,365 | 199,581 | 1,412,598 | 669,746 | 8,161,295 | 20,232,173 | 215,777 | 1,330,338 |
| DNA/Kolobok | 78,648 | 0 | 280,388 | 8,450 | 0 | 69,031 | 64,765 | 0 | 0 |

Supplementary Table 6: Support values for synteny analyses for Figure 3 and Supplementary Figure 7
*see separate excel file

Supplementary Table 7: The names used for chromosomes in the manuscript vs. their accession number/sequence name
*see separate .tsv file

Supplementary Table 8. Specimen data and accession numbers of all WGS *Nanomia* samples that are introduced in this study. CWD16 and NA19 are from Ahuja et al. 2024 but are included in the table.

| ID | sample_ID | collection_ID | ocean | location | lat_long | date_collected | depth |
|---|---|---|---|---|---|---|---|
| YPM-IZ-35039 | CWD16 | OC379-11-7 | NW Atlantic | Virginia | 37.450000, -74.016667 | 2002-06-08 | 0-20 m |
| YPM-IZ-111736 | WS5 | D325-SS12, #9 | NE Pacific | California | 36.600636, -122.376887 | 2011-12-02 | 2854.16 |
| YPM-IZ-111737 | WS6 | D326-SS1, #28 | NE Pacific | California | 36.071808, -122.282111 | 2011-12-03 | 2563.33 |
| YPM-IZ-111738 | WS7 | D331-SS5a, #136 | NE Pacific | California | 36.699365, -122.101074 | 2011-12-07 | 662.29 |
| YPM-IZ-111739 | WS8 | D500-SS12, #190 | NE Pacific | California | 36.647412, -122.113209 | 2013-07-15 | 559.83 |
| YPM-IZ-107008 | WS9 | D614-SS12, #225 | NE Pacific | California | 35.520, -122.5239 | 2014-05-22 | 309 |
| YPM-IZ-111745 | WS10 | V3557-D4, #51 | NE Pacific | California | 36.700917, -122.049541 | 2010-05-03 | 217 |
| YPM-IZ-111742 | NA31 | GOCBW2 nanomia | Gulf of California | Gulf of California | 27.591667, -111.516667 | 2010-06-18 | 0-20 m |
| YPM-IZ-111740 | NA32 | GoC2015 BW6 nanomia | Gulf of California | Gulf of California | 22.916667, -108.116667 | 2015-03-13 | 0-20 m |
| YPM-IZ-111741 | NA33 | GoCBW8 nanomia | Gulf of California | Gulf of California | 24.183333, -108.633333 | 2015-03-15 | 0-20 m |
| YPM-IZ-111733 | NA29 | KOK BW4-2 | N Pacific | Hawai'i | 19.344170, -156.075853 | 2017-03-16 | 0-20 m |
| YPM-IZ-110451 | NA28 | RI2-5 | NW Atlantic | Rhode Island | 41.036, -71.399 | 2013-08-30 | 0-20 m |
| YPM-IZ-110216 | WS2 | RI2-8 | NW Atlantic | Rhode Island | 41.036, -71.399 | 2013-08-30 | 0-20 m |
| YPM-IZ-111734 | WS3 | RI8-28 #24: | NW Atlantic | Rhode Island | 41.135, -71.366 | 2015-09-15 | 0-20 m |
| YPM-IZ-111735 | WS4 | RI6-12-C10 | NW Atlantic | Rhode Island | 41.056, -71.342 | 2014-08-20 | 0-20 m |

| | | | | | | | |
|---|---|---|---|---|---|---|---|
| YPM-IZ-111732 | NA30 | V13 | Mediterranean | France | 43.690000, 7.310000 | 2011-04-01 | 0-20 m |
| YPM-IZ-111164 | NA34 | Nanomia-FHL-A | NE Pacific | Washington | 48.54505, -123.012141 | 2024-06-16 | 0-20 m |
| YPM-IZ-111165 | NA35 | Nanomia-FHL-B | NE Pacific | Washington | 48.54505, -123.012141 | 2024-06-16 | 0-20 m |
| YPM-IZ-111166 | NA36 | Nanomia-FHL-C | NE Pacific | Washington | 48.54505, -123.012141 | 2024-06-16 | 0-20 m |
| YPM-IZ-111590 | NA37 | HI-NA-43; tube 1265 | N Pacific | Hawai'i | 19.6183, -156.0842 | 2024-04-12 | 0-20 m |
| YPM-IZ-111591 | NA38 | HI-NA-42; tube 1267 | N Pacific | Hawai'i | 19.6183, -156.0842 | 2024-04-12 | 0-20 m |
| YPM-IZ-111744 | NA19 | V3698-D2 | NE Pacific | California | 36.698983, -122.050126 | 2013-02-11 | 296 |
| YPM-IZ-111756 | | D1399-SS10 | NE Pacific | California | 36.697371, -122.071812 | 2021-10-30 | 447 |

Refs:


Cantalapiedra, Carlos P., Ana Hernández-Plaza, Ivica Letunic, Peer Bork, and Jaime Huerta-Cepas. 2021. “eggNOG-Mapper v2: Functional Annotation, Orthology Assignments, and Domain Prediction at the Metagenomic Scale.” *Molecular Biology and Evolution* 38 (12): 5825–29.

Condamine, Thomas, Muriel Jager, Lucas Leclère, Corinne Blugeon, Sophie Lemoine, Richard R. Copley, and Michaël Manuel. 2019. “Molecular Characterisation of a Cellular Conveyor Belt in Clytia Medusae.” *Developmental Biology* 456 (2): 212–25.

Holland, P. W., H. A. Booth, and E. A. Bruford. 2007. “Classification and Nomenclature of All Human Homeobox Genes.” *BMC Biology* 5 (October). https://doi.org/10.1186/1741-7007-5-47.

Huerta-Cepas, Jaime, Damian Szklarczyk, Davide Heller, Ana Hernández-Plaza, Sofia K. Forslund, Helen Cook, Daniel R. Mende, et al. 2019. “eggNOG 5.0: A Hierarchical, Functionally and Phylogenetically Annotated Orthology Resource Based on 5090 Organisms and 2502 Viruses.” *Nucleic Acids Research* 47 (D1): D309–14.

Khalturin, Konstantin, Chuya Shinzato, Maria Khalturina, Mayuko Hamada, Manabu Fujie, Ryo Koyanagi, Miyuki Kanda, et al. 2019. “Medusozoan Genomes Inform the Evolution of the Jellyfish Body Plan.” *Nature Ecology & Evolution* 3 (5): 811–22.

Schultz, Darrin T., Steven H. D. Haddock, Jessen V. Bredeson, Richard E. Green, Oleg Simakov, and Daniel S. Rokhsar. 2023. “Ancient Gene Linkages Support Ctenophores as Sister to Other Animals.” *Nature* 618 (7963): 110–17.

Simakov, Oleg, Jessen Bredeson, Kodiak Berkoff, Ferdinand Marletaz, Therese Mitros, Darrin T. Schultz, Brendan L. O’Connell, et al. 2022. “Deeply Conserved Synteny and the Evolution of Metazoan Chromosomes.” *Science Advances* 8 (5): eabi5884.
