## Supplementary Figure 1 for "A Chromosome-Scale Genome of *Nanomia septata* Reveals Extensive Rearrangement But No Clear Driver of the Unique Colony-Level Organization of Siphonophores"

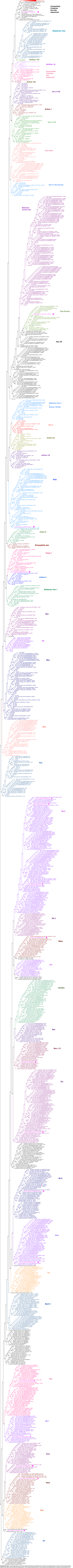

**Supplementary Figure 1**. Antp homeobox gene tree (Hox and Nk-related gene families). Maximum likelihood tree of 1,465 Hox and Nk-related sequences, built under model LG+R8, with ultrafast bootstrap branch support (1,000 replicates) values along branches. This tree contains protein sequences from the orthogroup with Antp homeobox genes, in addition to previously identified Hox/Hox-related sequences from Khalturin *et al.* (2019) and Steinworth *et al.* (2022). *Nanomia septata* sequences are colored in magenta and highlighted with a star. Sequence names retrieved from Antp orthogroup contain the species name (followed by _pep), the sequence ID from original source file and a label with the functional annotation generated by eggNOGG-mapper. Branch lengths were transformed to equal for display. Original tree is available in the supplementary data folder.
