## Supplementary figures and images for "A Chromosome-Scale Genome of *Nanomia septata* Reveals Extensive Rearrangement But No Clear Driver of the Unique Colony-Level Organization of Siphonophores"

### Supplementary Figure 2

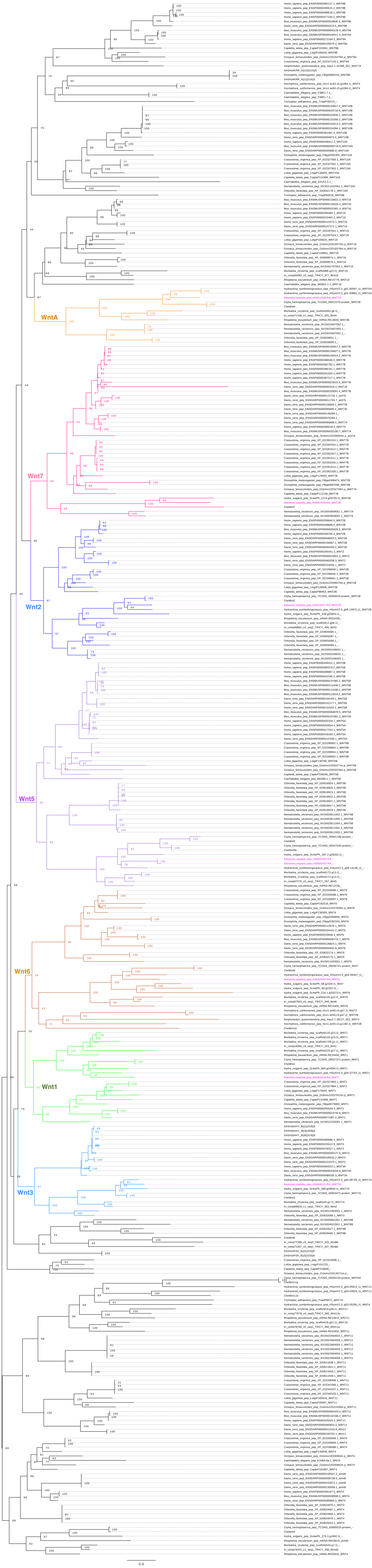
